# Widespread Occurrence and Diverse Origins of Polintoviruses Influence Lineage-specific Genome Dynamics in Stony Corals

**DOI:** 10.1101/2023.10.06.561300

**Authors:** Danae Stephens, Zahra Faghihi, Mohammad Moniruzzaman

## Abstract

Stony corals (Order *Scleractinia*) are central to vital marine habitats known as coral reefs. Numerous stressors in the Anthropocene are contributing to the ongoing decline in coral reef health and coverage. While viruses are established modulators of marine microbial dynamics, their interactions within the coral holobiont and impact on coral health and physiology remain unclear. To address this key knowledge gap, we investigated diverse stony coral genomes for ‘endogenous’ viruses. Our study uncovered a remarkable number of integrated viral elements recognized as ‘Polintoviruses’ (Class *Polintoviricetes*) in 30 *Scleractinia* genomes, with several species harboring hundreds to thousands of polintoviruses. We reveal massive paralogous expansion of polintoviruses in stony corals, alongside presence of integrated elements closely related to Polinton-like viruses (PLVs), a group of viruses that exist as free virions. These results suggest multiple integrations of polintoviruses and PLV-relatives, followed by their paralogous expansions shaped stony coral genomes. Gene expression analysis reveals all polintovirus structural and non-structural hallmark genes are expressed, strongly supporting free virion production from polintoviruses. Our results revealing a significant polintovirus diversity across the *Scleractinia* order open a new research avenue into their possible roles in disease, genomic plasticity, and environmental adaptation in this key group of organisms.

## Introduction

Coral Reefs are a striking feature of many marine habitats, glowing with an abundance of life of every size, shape and color. These reefs are formed by a group of clonal composite symbiotic organisms (order *Scleractinia*) with individual colonies composed of genetically identical polyps hosting single celled algae of the *Symbiodiniaceae* family, working together to secrete and cement calcium carbonate skeletons into vast three-dimensional structures ^1–3^. Globally, coral reefs have been estimated to harbor 25% of marine biodiversity ^4^ and even beyond that scope, through recreation, tourism, and natural resource use, they are estimated to provide around $2.7 trillion USD worldwide in economic value each year ^5^. However, for as much as we value these reefs and their socioeconomic contributions, coral reef health and coverage are in decline, with coverage dropping from 33.3% in 2009 to 28.8% in 2018 ^5^, with that declining trend only accelerating in recent years.

There are multiple factors impacting this decline, which include warming oceanic temperatures (inducing longer lasting, higher intensity, and more frequent long term mass bleaching events and their associated mortality ^5–9^), decreased pH (and the associated lowered calcification rate slowing colony growth^5, 6^), and the rapid spread and transport of coral diseases across colonies and between reef sites.^5, 9, 10^. These factors have different levels of impact on coral reefs across the world, accounting for the main stressors in any given local environment. For example, in the Caribbean specifically, it is disease that plays the largest role in coral mortality and reef decline^11^. Of the great number of disease outbreaks occurring across corals over the years^12–14^ the most well-known and devastating outbreaks are those that occurred in the Caribbean^11, 15–18^. These various diseases have originated due to a number of factors, with many diseases having multiple compounding causal agents^12, 13^, viruses being one of them.

Virus-host interactions in coral have a number of effects, from the more traditional role of viruses as agents of infection and disease^19–22^, to alterations of the host holobiont and its associated communities impacting host health and resilience ^20, 22^. In the context of coral holobionts, the role of viruses could be both beneficial and harmful. These effects potentially encompass pathogenic interactions, as well as important top-down control mechanisms that constrain the microbiome community dynamics ^23, 24^, in addition to interactions that potentially result in increased immunity and protection from other pathogens^20, 23, 25–27^.

So far, multiple approaches have been used to examine viral interactions with coral, from transmission electron microscopy in early reports examining the presence of virus like particles within coral tissue^28, 29^, to metagenomic approaches examining both coral and symbiont strains in diverse locations and under diverse environmental conditions^4, 30–32^. Despite this, there has been little work done to determine the function and identity of most members of the holobiont virus communities within coral^32^, which remains an important task given the key contribution of viruses in diverse ecosystems.

One underexplored aspect of virus-host interactions with potential implications for coral health and holobiont stability is the level of viral endogenization within coral host genomes. Endogenization refers to the integration of near-complete or partial viral genomes into the host genome. Endogenous viruses exist in varying levels within the genomes of numerous eukaryotes^33–35^, and the process of endogenization has been ongoing for millions of years throughout the course of eukaryotic evolution^24, 36–38^. These elements can be incredibly diverse, with instances of hundreds to thousands of such elements inhabiting a single genome^25, 39^, and large-scale endogenization of near-complete genomes of ‘giant viruses’, comprising a high percentage of the total genome content in many eukaryotic species^33–35, 40^.

Endogenous viruses reveal the history of past interactions of hosts with viruses, as the endogenization process leaves traces of past infections in the genomes of their hosts^23, 24, 40, 41^. Thus, endogenous viral elements (EVEs) can provide an opportunity to reveal the phylogenetic identity of extant viruses that infect a particular host lineage. A notable, relevant study in this regard is the identification of RNA viruses infecting the Symbiodiniaceae members of coral holobionts based on RNA virus EVE signatures in *Symbiodinium* genomes^42^.

Motivated by the lack of information on the interaction of viruses with corals, we performed a comprehensive investigation of the stony coral genomes for endogenous virus signatures with a goal to shed light on the roles of such elements in coral genome evolution along with identifying the viral lineages that likely infect corals in nature. Intriguingly, our study led to the identification of ‘Polintoviruses’ residing within the genomes of stony, reef building coral (Order *Scleractinia*) from 11 different genera. Polintoviruses, also known as Maverick/Polinton elements, were first reported in 2005^43^. Originally defined as transposable elements (selfish genetic elements that can proliferate within the genome of the host^44–47^) these are large elements approximately 10kb to 50kb in size^48, 49^, representing some of the largest integrating DNA-elements known. Later discovery of conserved structural proteins (major and minor capsid proteins; MCP and mCP respectively) in these elements prompted re-classification of these elements as viruses^50^, as capsid proteins are considered to be a key and defining characteristic of viruses ^51^. This classification of polintoviruses as *bona fide* viruses was officially recognized by International Committee on Taxonomy of Viruses (ICTV) - as class Polintoviricetes was created within which a single order Adintoviridae include these elements^50, 52^.

Further study of the MCP of polintoviruses determined it to be closely related to the MCPs of virophage^53^; a class of small dsDNA viruses that need Nucleocytoplasmic large DNA viruses (NCLDVs) for replication^54^. This led to the hypothesis that virophages originated from Polintoviruses^50, 52^. This enhanced understanding led to the discovery of Polinton-like viruses (PLVs) in 2015^53^. PLVs as well as a few close relatives of PLVs that harbor polintovirus-like characteristics were found to be highly abundant in aquatic environments^48^.

In this study, we report the widespread prevalence of polintoviruses and relatives of PLVs within diverse stony coral genomes, and analyze their phylogenetic histories, gene expression dynamics and functional potential. Our study opens the door for investigations into the origins of these elements and their impact on coral physiology, environmental adaptation, and genome evolution.

## Results and Discussion

### Prevalence of polintoviruses in stony coral genomes

Using the bioinformatic pipeline we established for screening viral regions originating from polintoviruses (See Methods), polinton-like viruses (PLVs) and virophages, we identified numerous endogenous viral regions (EVRs; Fig.1.B) within each of the thirty analyzed *Scleretina* genomes (Genome accession numbers in Supplementary Table 1). Our preliminary screening utilized a strict cut-off criterion dictating that each region needed to contain a major capsid protein (MCP), minor capsid protein (mCP), and ATPase gene - genes that are commonly present and conserved across the members of class *Polintoviricetes*, virophages and polinton-like viruses (PLVs)^48, 51^. These EVRs varied in length widely within our specified size cut-off (10kb-50kb; Supplementary Fig.1) consistent with what has been previously found in several studies^39, 55^. The overall mean and median length for validated EVRs were found to be 23,555 bp and 21,932 bp, respectively, with an overall right-skewed distribution of EVR lengths (Supplementary Fig.2).

**Figure 1:**
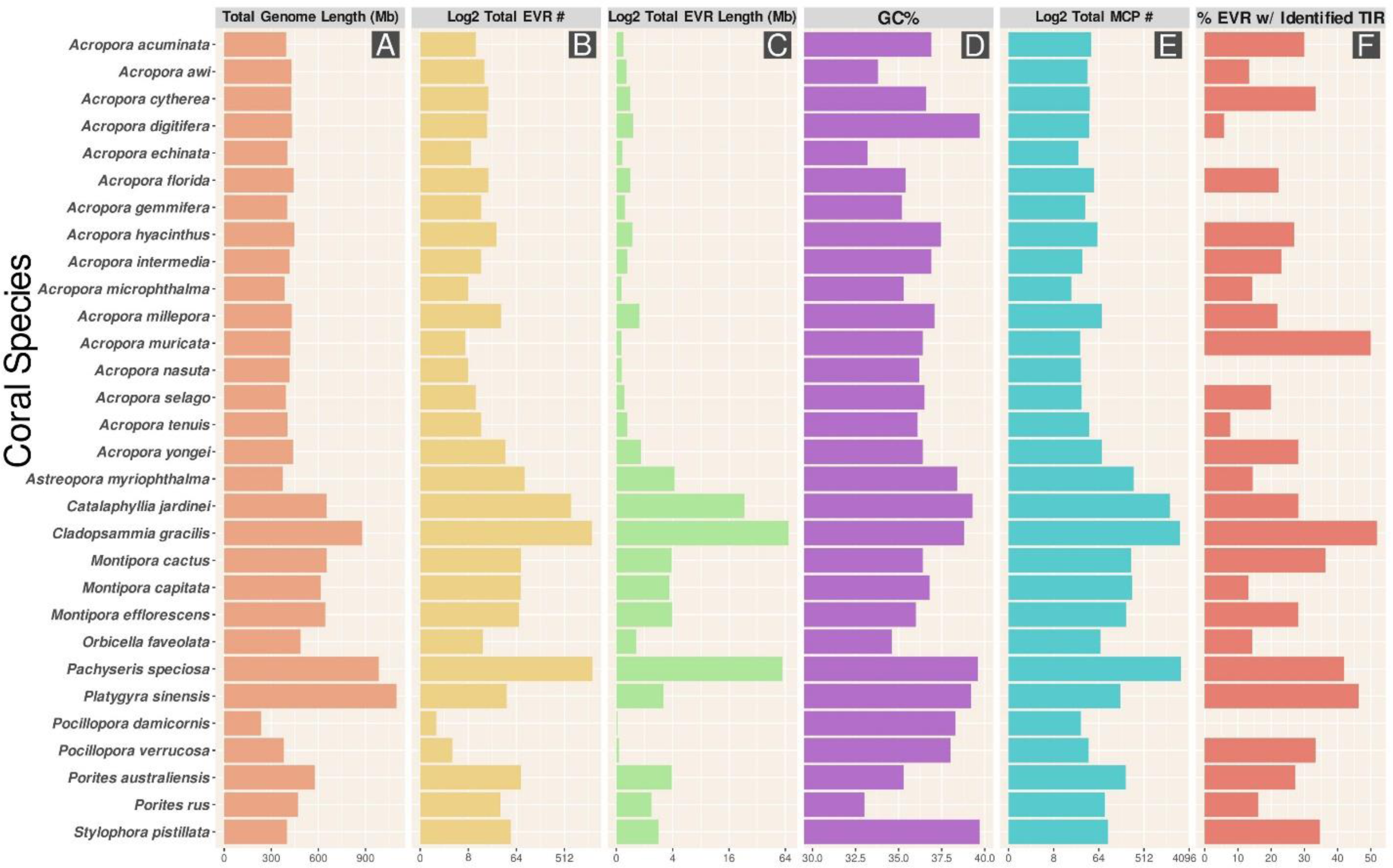
Bar Plots of Coral Species in regards to **(A)** Total Genome Length, measured in Million Base Pairs (Mb), **(B)** Total Number of endogenized viral regions (EVR), represented on log2 scale, **(C)** Combined Length of all EVRs within a genome measured in Megabase pairs (Mb) and represented on a log2 scale, **(D)** Overall percentage of GC content of EVRs in each coral genome, **(E)** Total Number of Identified MCP genes per coral genome, represented on log2 scale, **(F)** Proportion of EVRs that harbor Terminal Inverted Repeats.

When we utilized a more relaxed quality control criteria (set to include any three hallmark genes, regardless of type), we found that on average 26.78% more putative EVRs passed cut-off requirements (Paired t-test, p-value = 0.0212). This indicates that our results represent a conservative estimate of polintoviruses in stony coral genomes. Due to the understudied nature of polintoviruses, it is likely that substantial sequence diversity exists in some of their core genes that render them difficult to be accurately identified with existing information on their hallmark genes, such as the highly divergent MCP homologs identified by Bellas *et al.* (2023)^39^. We chose to use stringent quality controls to ensure the validity of our initial exploratory results, however we note that many of the EVRs we excluded may in fact be *bona fide* viral candidates.

In addition to identifying the viral regions, we also searched specifically for MCP homologs (within or outside the viral regions) across the genomes, and we found consistently higher numbers of MCPs in most genomes than validated EVRs (Fig.1 B,E). We found that number of MCPs in the stony coral genomes were on average 34.7% more than the number of EVRs, with the largest discrepancy between these counts observed in *Cladopsammia gracilis (C. gracilis)* (62.4% more MCP hits than EVR counts) and the lowest *Pocillopora damicornis (P. damicornis)* genome (3.6%). This supports the notion that the ‘true’ number of polintovirus EVR in stony coral species are likely higher than what we have found. This observation is also supported by Bellas *et al* (2023)^39^, who found that the integrated MCPs are usually part of endogenous viral elements, which might not always be detectable due to low quality genome assemblies from short reads.

This discrepancy between the number of EVR and MCP candidates recovered may be due in part to the fact that the coral genomes analyzed in our study used varying sequencing and assembly approaches. It has been demonstrated that short read sequencing techniques, such as Illumina^56^, are limited in their ability to capture the entirety of endogenous viruses, due to fragmented assemblies of eukaryotic genomes, which are often full of repetitive elements, along with miss-assemblies, thus introducing frameshifts or other errors that make identification of such regions difficult. In addition, fragmented assemblies often lead to small contigs which might contain one or two viral hallmark genes but will not represent the entire EVR. This possibility is supported by a recent study by Bellas *et al*. (2023)^39^, who have reported recovery of complete endogenized PLVs in draft genomes after reassembly and error-correction of several protist genomes using long-read sequencing technology.

In further support of this, an additional study by Noel *et al* (2023)^57^ determined that long read sequences were essential for developing and obtaining an accurate understanding of gene content, as their work in assembling coral genomes from Tara Pacific^58^ data demonstrated that a great deal of ‘repeat’ and duplicated genetic elements were lost using short read data. Taken together with the findings of Bellas *et al*. (2023)^39^, it is highly likely that contig fragmentation, gene loss, or divergent copies of core genes are responsible for these ‘missing’ core genes in our identified EVR. Nevertheless, we found a strong positive correlation between the number of EVR (Fig.1 B) and the number of MCP hits (Fig.1 E) across genomes, (Spearman’s ρ = 0.947), indicating that while the number of EVR we identified might be conservative, it is nevertheless a good indicator of the genus or species-specific variations in polintovirus abundances across the stony coral genomes.

We also sought to identify the presence of Terminal Inverted Repeats (TIRs) within our EVRs. TIRs are hallmarks of inserted elements^59^, and previous studies have found that in protist genomes, as much as 50% of the polintoviruses located on long contigs can harbor TIRs at their ends^39^. We found that many of our identified EVR candidates harbor TIRs, with the highest percentage of validated EVRs containing TIRs was found within *C. gracilis*; wherein 869 of the 1676 validated EVRs possessed TIRs (Fig.1 F; Supplementary Fig.3). On average 22.8% of the identified EVR across the coral genomes harbored TIRs. It is likely that we were unable to detect additional TIRs-containing EVRs given the genome assembly quality - with many of these regions residing on small contigs.

Overall, some of the stony coral genomes harbored a remarkably high number of EVRs and MCPs. Of note are the genomes of *C. gracilis* and *Pachyseris speciosa (P. speciosa),* with the former harboring 1,676 EVRs and 2,686 MCPs, and the latter harboring 1,724 EVRs and 2,802 MCPs. All other genomes examined contained less than 100 EVRs and 320 MCP hits; except for *Catalaphyllia jardinei (C. jardinei)*, which harbored 680 EVRs and 1,685 MCP hits. The genome with the least number of hits was *P. damicornis*, harboring only 1 EVR that we were able to identify within our specified criteria (Fig.1.B; Supplementary Fig.4). These results reveal the remarkable variations in prevalence of polintoviruses across different members of the Scleractinia order.

The smallest of the genomes we analyzed was *P. damicornis* (234.35 Mbp), and the largest was *Platygyra sinensis* (*P. sinensis*) (1096.36 Mbp) (Fig.1.A; Supplementary Fig.6). We identified a positive correlation between length of coral genome and number of EVR (Spearman’s ρ= 0.73) (Supplementary Fig.7). We likewise found a positive correlation between the length of the coral genome and the number of MCP hits (Fig.1.E; Supplementary Fig.8; Spearman’s ρ = 0.69), indicating that at least in the limited subset of genomes we analyzed, longer genomes appear to usually harbor larger numbers of EVRs. Interestingly we found that *P. sinensis*, despite having the longest genome (at 1,096 Mbp), contained only 41 EVR and 172 MCP hits, whereas *P. speciosa* at the second longest (984 Mbp) held the second highest number of EVRs (Fig.1.A,B,E). This observation suggests that factors other than genome sizes are also responsible for the lineage-specific variations in the EVR abundances we observed.

Due to these vast numbers of EVRs, in several genomes they contributed a sizable portion of these genomes (Fig.1.C) - for example ∼7.8% of the *C. gracilis* genome is composed of these EVRs (Supplementary Fig.5). Indeed, if we consider the possibility that most of the MCP hits in this genome are part of endogenous polintoviruses, then the true contribution of polintoviruses to the genome content of *C. gracilis* will be even higher. Although relatively rare, similar massive proliferation of polintoviruses has been reported in protist *Tritrichomonas vaginalis,* where approx. 27-54% of the genome length was contributed by these elements (Bellas *et a*l. 2023). The same study also reported that ∼10% of the genome of *Paulinella micropora* consists of PLVs. As far as we are aware, *P. speciosa* has the largest number of polintovirus regions ever reported in metazoans.

### Phylogeny of the stony coral EVRs

For phylogenetic analysis, we first clustered the MCPs from the identified EVRs at a 90% amino acid sequence similarity to reduce redundancy in the dataset (See Methods). In doing so we determined that the number of MCP clusters are quite high in most genomes, indicating high within-genome diversity of polintoviruses in stony corals. A similar level of diversity of PLVs were reported in several protist genomes, indicating the possibility of multiple independent integration events shaping the diversity within these genomes^39^. We then performed a maximum likelihood phylogenetic assessment on the representative sequences from each of these clusters along with reference Adintoviridae members, PLV and virophage MCPs (See Methods) to clarify the evolutionary history of the EVRs in stony corals. (Fig.2; Supplementary Fig.9)

**Figure 2:**
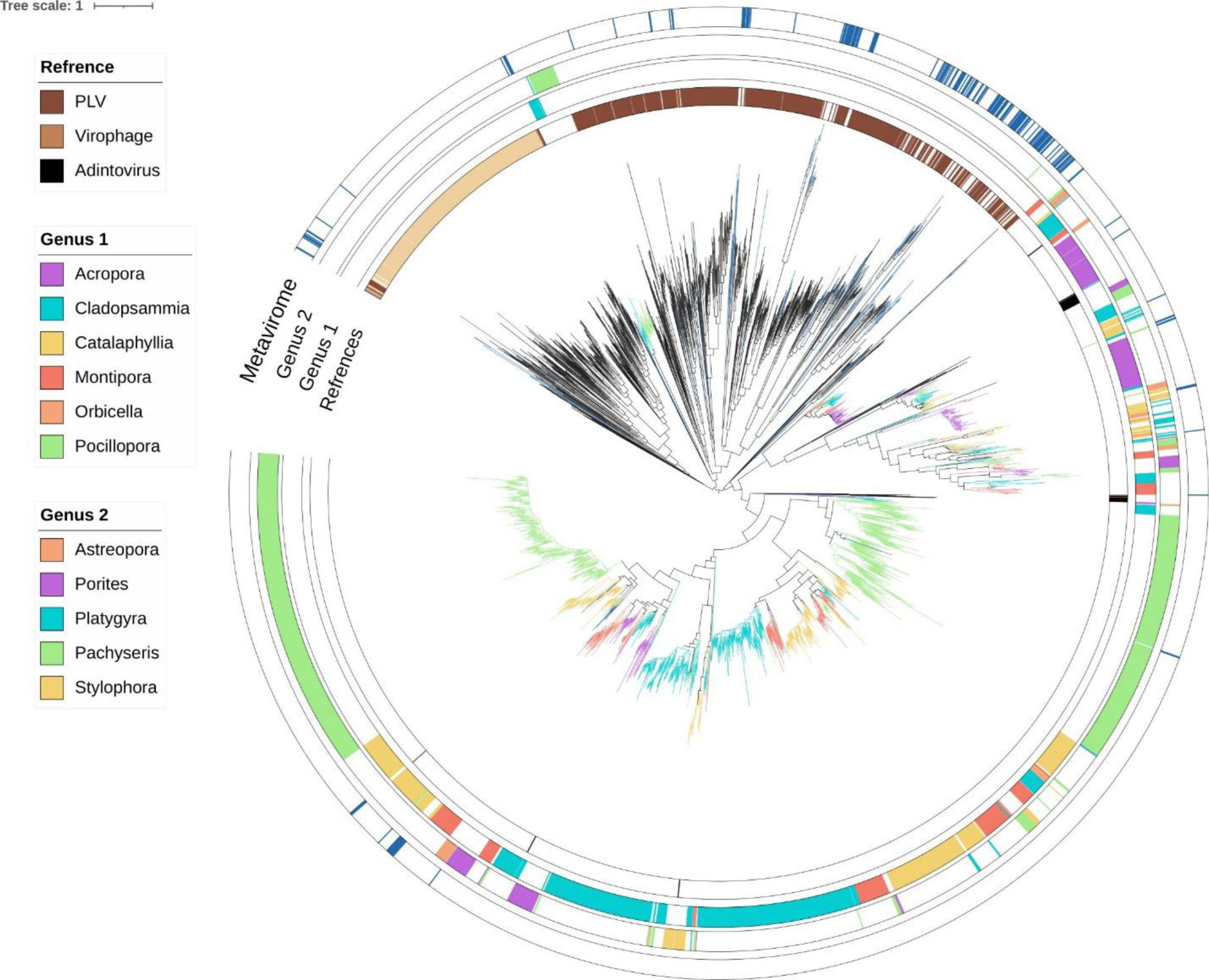
Maximum likelihood phylogeny of major capsid proteins (MCPs) from EVRs in stony coral genomes, stony coral viromes and reference sequences belonging to *Adintoviridae* members, polinton-like viruses (PLVs) and virophages. **Ring 1 (Intermost)**: Reference sequences. **Ring 2 and 3:** EVRs in Stony Coral (By Genus). **Ring 4 (Outermost):** Viruses recovered from stony coral viromes.

We found that the vast majority of the stony coral EVRs clustered separately from the environmental PLVs and virophages (Fig.2), supporting previous observations^51, 60^ regarding the evolutionary history of these elements. Most of the environmental virophages and PLVs formed their own distinct clusters on the phylogeny, although a few of the PLVs did cluster with the virophages. We used a recently proposed homology-based approach to identify the virophages in our datasets^61^(See Methods), and it is possible that a few false positives emerged in this classification approach. In contrast, the majority of the stony coral polintoviruses formed separate clusters along with the members of the previously established *Adintoviridae* family. Adintoviruses were reported within diverse animal genomes, which have since been identified to be essentially polintoviruses, and thus were later classified within the class *Polintoviricetes*^62^.

Polintoviruses within the stony coral genomes formed two large monophyletic clusters, each cluster harboring sequences from most of the genus, suggesting presence of phylogenetically distinct polintoviruses within the same lineage (genus or species) of stony coral (Fig.2); consistent with multiple origin of these elements throughout the evolutionary history of the Scleractinia order. Phylogenetic analysis showed distinct patterns of paralogous expansion of stony coral polintovirus genomes. Specifically, a large clade within the tree where the majority of the EVRs are clustered (except *Acropora*), showed large expansion of these elements. The largest degree of such expansion was found to be in the species *C. gracilis*, *C. jardenei*, and *P. speciosa*, consistent with the fact these three species harbored the largest number of these elements overall (Fig.1.B). A separate cluster of stony coral polintoviruses that also harbored representatives from most genus did not show as much paralogous expansion compared to the cluster previously discussed. All the EVRs from different *Acropora* species are also clustered within this group. We observed a similar phenomenon of genus-specific clustering of EVRs for other coral genomes for which we were able to analyze multiple species; specifically, *Monitopora* (n=3), *Porites* (n=2) and *Pocillopora* (n=2). While EVRs from the same genus generally cluster together, we also found several instances where EVRs from multiple genus clustered within close proximity to each other, suggesting exchange of these elements across different genera.

To date, a comprehensive assessment of the evolutionary origin of polintoviruses remains to be performed, despite their identification in a number of vertebrate and invertebrate genomic lineages^33–35^. Our observation of distinct patterns of expansion of the stony coral polintoviruses shaped by their phylogenetic history suggests that while some of these elements went through within-genome amplification upon integration, others were not subject to such expansion to a similar degree. Paralogous expansion of endogenized polintoviruses constrained by the phylogenetic history might provide some type of fitness benefit to the host - a possibility that needs further scrutiny. Future studies focusing on the molecular underpinnings of the proliferation dynamics of these elements could address questions regarding the genomic and evolutionary factors shaping the dynamic expansion landscape of polintoviruses in stony corals.

Even though the vast majority of the stony coral EVRs clustered distinctly from the environmental PLVs, we observed clustering of several EVRs with the established phylogenetic group of environmental PLVs (Fig.2). Specifically, a number of the elements from the *Cadopspammia, Pachyserys*, and *Platygyra* genera clustered within the monophyletic group of PLVs. As discussed previously, *Cladopspammia* and *Platygyra* also harbor a large number of EVRs in general, along with high paralogous expansion. The environmental PLV sequences in our tree originated from free virus particles^39^, thus the clustering of some of the stony coral EVRs within this clade supports the possibility that along with polintoviruses, viruses having close evolutionary ties with PLVs potentially infect and integrate within stony coral genomes.

Results from the phylogeny paint a highly dynamic genomic landscape of the stony coral polintoviruses with multiple distinct origins and large-scale lineage-specific expansion likely constrained by their evolutionary histories. The varying degree of expansion of these elements in different stony coral genomes and possible horizontal transfer between genus makes them a key player in the genome evolution of stony corals. Transfer of transposable elements across closely related lineages is not uncommon^63^. Consistent with our observations, an analysis from a very recent study by Jeong *et al*. (2023)^64^ found the horizontal transfer of polintoviruses across different nematode strains within the same species, and even inter-phylum transfer of these elements. Presence of both phylogenetically distinct polintoviruses and close relatives of PLVs across different stony coral species is consistent with ancestral integration of diverse polintoviruses and PLV-like viruses within host genomes, followed by vertical transfer and lineage-specific proliferation. Further exploration of additional coral genomes as well as a more comprehensive phylogeny will better clarify the evolutionary history of endogenous coral viruses.

### Prevalence of polinton-like viruses in the virome of stony corals

The abundant and widespread presence of polintoviruses and close relatives of PLVs we uncovered within *Scleractinia* genomes warranted further investigation on whether free viruses, closely associated with polintoviruses or PLVs, could be part of the coral virome. To accomplish this, we leveraged the raw virome sequence data from a study conducted by Weynberg *et al*. in 2017^32^. The authors of this study performed a CsCl centrifugation method on several environmental stony coral species to isolate virus-like particles with minimal host contamination to establish the overall viral community. CsCl centrifugation is a widespread technique used in environmental virome studies to isolate virus-like particles^65^. This circumvented the methodological limitation of distinguishing virus particles from the coral tissue samples during isolation and sequencing of these components.

Searching this dataset using the MCP HMMs we constructed (See Methods) revealed a total of 371 hits to MCP proteins in the virome of several stony coral species. After incorporating these MCPs (following de-replication at >95% amino acid identity) within our existing MCP phylogenetic framework, we found that vast majority of these sequences cluster within the environmental PLV clade in our tree, while a few of them were closely affiliated with the virophages (Fig.2). Interestingly, some of these viral sequences are also clustered within the large clades of stony coral polintoviruses; thus, suggesting that free viruses of high similarity to the polintoviruses could be present in coral viromes. While the possibility of some host contamination cannot be completely ruled out, we do note that the estimated host contamination in these libraries is quite minimal at ≤1.5%^32^.

There was not much overlap between the species we analyzed for the EVRs and those sampled for virome by Weynberg *et al*. (2017) ^32^, with only three species, *P. damicornis, A. tenuis, and P. verrucosa* being common between these two datasets. Despite this limitation, results from the virome revealed PLVs to be an integral component of the stony coral virome, with the potential to infect eukaryotic members of the holobiont, including the coral animal. A targeted approach will be needed in future studies to definitely identify the potential hosts, as well as understand their impact on reef microbial dynamics and impacts on the health and resilience of stony corals themselves.

### Functional profile of the stony coral polintoviruses

To understand the functional diversity encoded within the polintovirus EVRs, we clustered the proteins based on sequence similarity and annotated representative proteins from each cluster. We included all the EVRs that were identified using both stringent and relaxed criteria (See Methods) to explore the full range of functional potential in these regions (Fig.3.A). Apart from the six known core genes that were highly prevalent in these regions as expected, we found several proteins with unknown function to be present in high frequency (>40% of the regions) in stony coral polintoviruses; including a protein homologous to poxvirus protein AMEV120, which contains a Pfam domain (DUF5679) of unknown function. Interestingly, in our analysis, another group of proteins emerged that are present on average in >10% EVRs, which included homologs of putative antirepressor protein YoqD, peptidoglycan binding domain, viral structural proteins and several proteins with unknown function.

**Figure 3:**
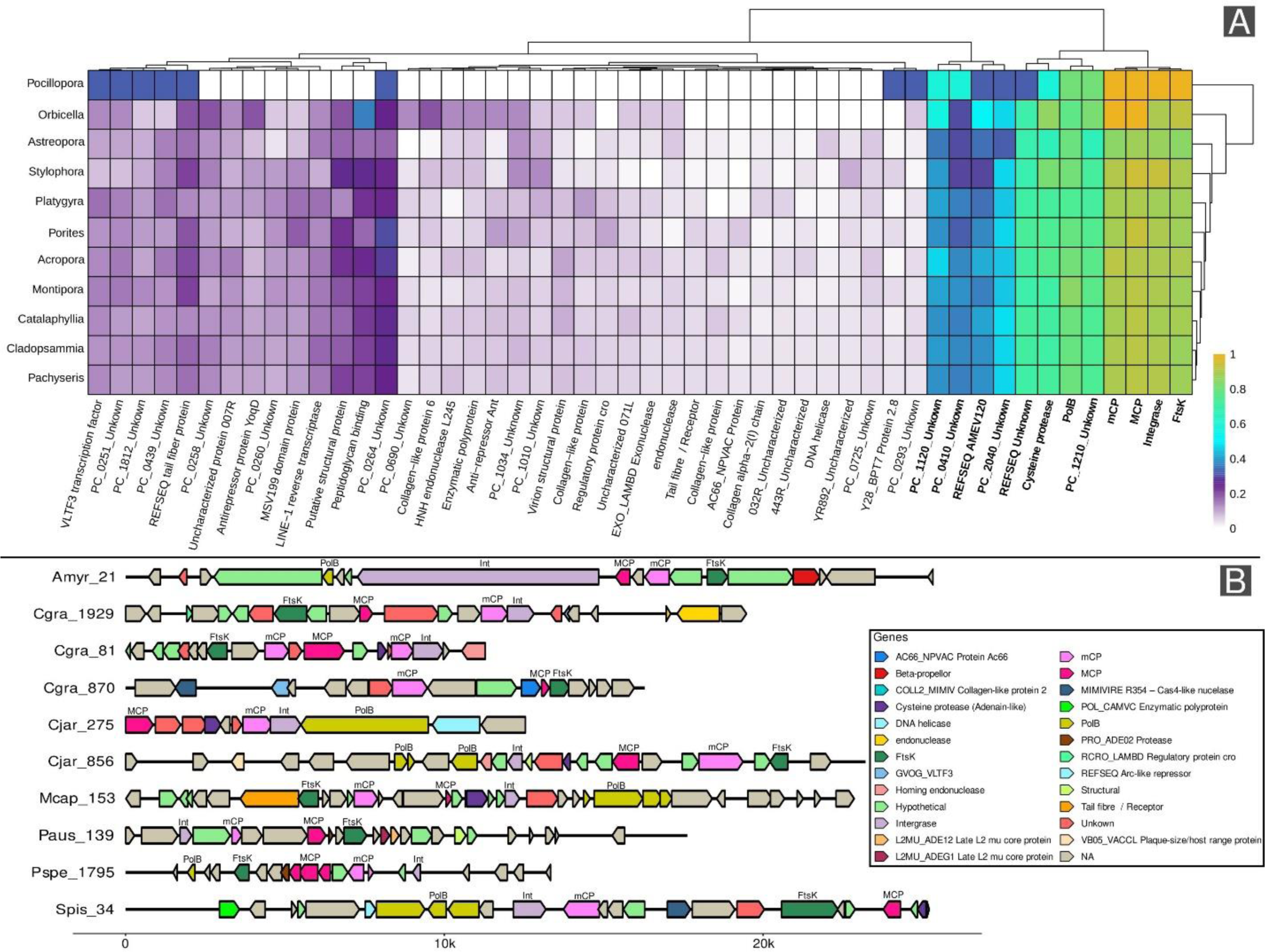
**(A)** Heatmap of relative proportion of different functions encoded by polintoviruses across different genus of stony corals. Each column represents the functional profile of polintoviruses in a particular genus. Hallmark (core) gene labels rendered in bold print. Annotated proteins occurring in at least 2% of total hits were clustered by coral genus and plotted. Heatmap was created in the ‘pheatmap’ R package ^101^. **(B)** Genome structure and gene locations of selected stony coral EVRs. Core genes labeled directly on the map. A selection of EVRs was chosen based on similar length and presence of notable genes (such as VLTF3). Figure was constructed using gggenomes package in R ^102^.

Also of interest, one of these proteins was homologous to the VLTF3 transcription factor, a gene known to be conserved in nucleocytoplasmic large DNA viruses (also known as giant viruses^40^; that infect diverse eukaryotes. The presence of this gene in a subgroup of polintoviruses have been reported before, with suggestions that exchange of this gene occurred between giant viruses and polintoviruses^66^. The presence of homologs to antirepressors and additional viral structural proteins further supports the fact that polintoviruses are *bona fide* viruses. However, much is yet to be discovered regarding the role of additional genes in dictating the life-strategies and evolutionary trajectories of polintoviruses. Finally, we found numerous proteins with low frequency (present in >3% of the elements) across the stony coral polintoviruses. Some of these are homologous to antirepressor proteins, helicases, virus structural proteins, exo- and endonucleases, and collagen-like proteins (Supplementary Fig.10).

This analysis revealed highly diverse functional profiles encoded across the stony coral polintoviruses, and the patchy presence of many of these genes across these viruses (Fig.3 B) suggest that their genomes (apart from the conserved core genes) were shaped by processes of gene gain and loss. This opens the question on the role of these genes in polintovirus propagation and their impact on host physiology and genome evolution. Recent studies have started to illuminate the accessory gene functions and factors shaping the genomic landscape of polintoviruses. Of note is a recent report on polintoviruses in nematodes, where these elements were found to be a vector of horizontal transfer of key nematode protease and kinase genes^67^. Specifically, polintoviruses took up these genes as cargoes and transferred them between different nematode species on a global scale. This presents evidence for the potential role of polintoviruses as agents of host genome evolution through the facilitation of horizontal gene transfer and the introduction of novel genetic elements.

### Polintovirus gene expression dynamics in *Stylophora pistillata*

Multiple studies have suggested that polintoviruses likely propagate as free virus particles, given the genomic evidence of presence of structural proteins (MCP and mCP)^48, 51, 66^. Yet, a clear understanding of the expression dynamics of the genes encoded in these viruses is lacking. In many cases, presence of a gene is not the direct evidence of its function, as gene expression of mobile genetic elements can be suppressed by host molecular machinery^68, 69^. Although previous studies have managed to identify the expression of polinton-like virus MCP from several protist genomes^39, 70^, a comprehensive assessment of the expression dynamics of the key genes in endogenous polintoviruses remain to be performed.

For an in-depth investigation of the expression patterns of stony coral polintoviruses, we leveraged the data by Rädecker N *et al*. (2021) ^71^ who sought to identify the transcriptomic response of *S. pistillata* to heat stress utilizing five genetically distinct coral colonies. We identified 49 polintovirus EVRs in the *S. pistillata* genome, all of which are distinct at 90% nucleotide identity. By mapping the transcriptomic data from Rädecker N *et al*. (2021) ^71^ to these elements, we are able to reveal several key aspects of the expression dynamics of polintoviruses. We found that the hallmark genes including MCP, mCP, ATPase, Integrase and PolB had detectable levels of expression across the samples in both heat stress treated and control samples (Fig.4.A). These genes were expressed in roughly 50% of the 49 polintoviruses (MCP = 40%, mCP = 41.67%, ATP = 39.13%, Integrase = 50.75%. PolB = 60.58%) (Supplementary Fig. 11) across different conditions. Because data in this report were generated with genets instead of replicated samples from an individual colony, statistical comparisons on overall levels of polintovirus expression in stressed vs. unstressed states cannot be made, but this analysis does provide evidence that all the key genes in these viruses are, in fact, being expressed (Fig.4.B).

**Figure 4:**
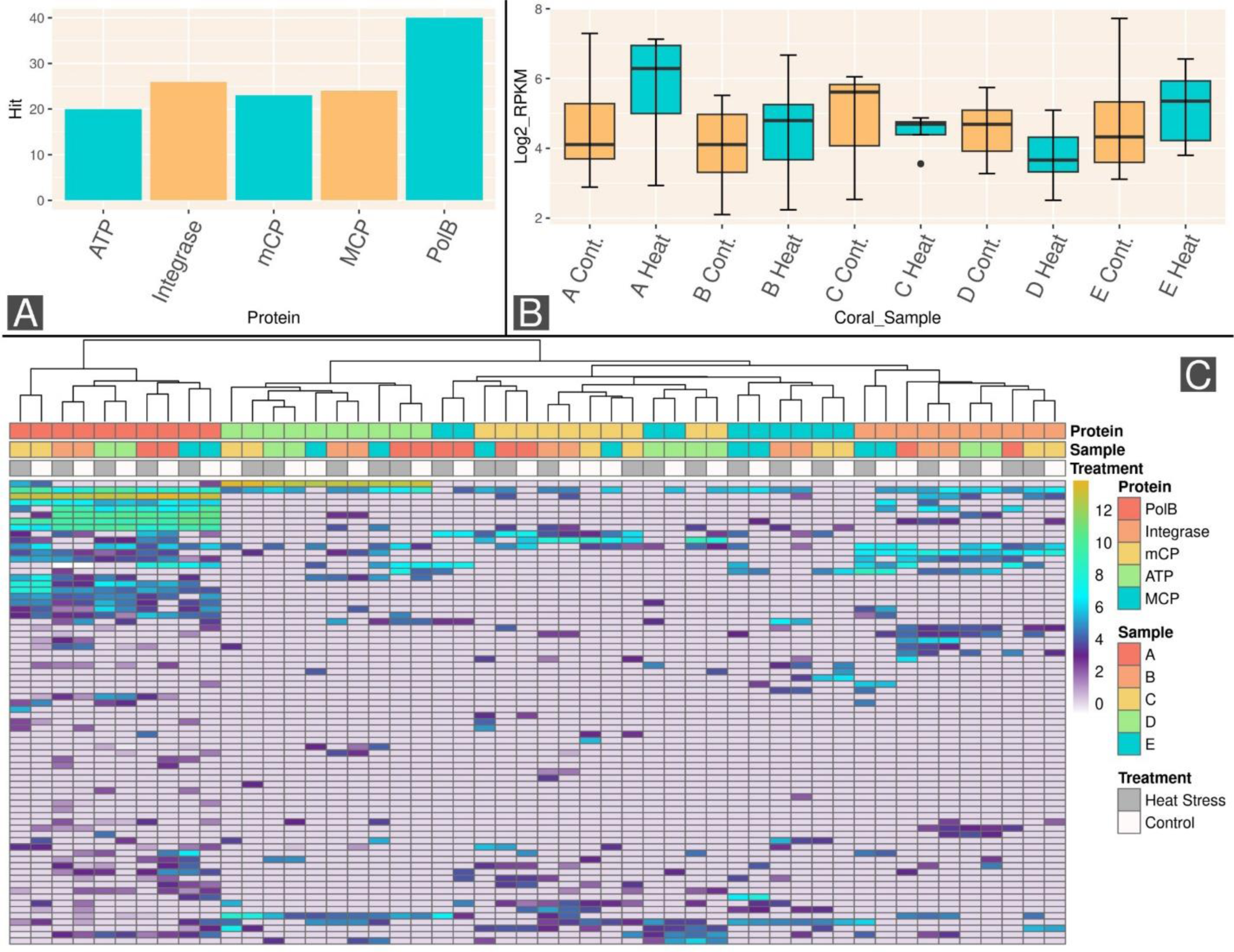
**(A)** Bar Plot of total *S. pistillata* viral regions expressing polintovirus hallmark Genes. **(C)** Boxplot of RPKM for detectable (RPKM >1) MCP expression in *S. pistillata* coral samples. **(D)** Heatmap of polintovirus core gene expression (PolB, ATPase, Integrase, MCP, and mCP) from *S. pistillata* viral regions. Expression levels shown in log2 scale. Annotation Layer 1 (Top): Protein identity. Annotation layer 2: Coral sample identity. Annotation layer 3 (bottom): Sample treatment type. Figure 4 C was plotted using the pheatmap ^101^package in R.

Additionally, hierarchical clustering of the hallmark genes expression levels revealed that the expression profile of individual genes across all polintoviruses primarily clustered by gene function (Fig.4.C). Data from PolB, ATPase and Integrase created gene specific clusters, whereas the structural proteins (mCP and MCP) formed their own unique cluster; separated from the other genes. This suggests that the transcriptional program of individual polintovirus genes is conserved across coral colonies. If the transcription of the key genes were ‘random’ or result of leaky transcription without an underlying molecular program, they would be expected to cluster randomly irrespective of gene functions, which is clearly not the case we observed.

Within each of these gene clusters, further subgrouping was driven primarily by samples, not by treatments (Fig.4 C). That is, data from individual colonies clustered together irrespective of stress treatments. Thus, expression of the individual polintovirus genes across different colonies show colony-specific differences. This phenomenon is similar to well-known biological differences in expression level of the same genes even in different strains of the same microbial species^72^. Therefore, any future studies wishing to examine differences in these expression levels due to various forms of stressors including those relevant to changing ocean conditions (temperature and pH, for example) should utilize clonal/fragmental replicates to avoid confounding effects introduced by biological or environmental variations among colonies of the same coral species.

The first study reporting polintoviruses, performed by Rinke *et al.* in 2006^43^, proposed a scenario of how polintoviruses propagate within the host genome. In this scenario, the polintovirus element is first excised from the host genome by the enzyme integrase, leading to an extrachromosomal single-stranded element, which is subsequently replicated by the PolB. After replication, the integrase enzyme catalyzes its integration in the host genome. However, this scenario does not take into account the function of other genes in the polintovirus genome. By utilizing the transcriptomic data, we were able to show that polintoviruses do express the genes involved in capsid formation (MCP and mCP) and genome packaging (ATPase; Fig.4; Supplementary Fig.12-16). These genes are only utilized in free living viruses, and thus would likely be lost during genome reduction in case of a strictly endogenous lifestyle. Conservation of these genes and their transcriptomic program supports the possibility that these elements re-enter the environment as free virions as part of their propagation. These scenarios are not mutually exclusive - it is possible that polintoviruses have both life-strategies that include within genome propagation and free particle production. A recent study also supports this possibility where it was shown that polintoviruses have horizontally transferred key genes across nematode lineages through a mechanism that requires free virus production^67^.

Our analysis of the stony coral viromes (discussed in a previous section) also revealed the presence of Polintovirus MCPs within the virome (Fig. 2). In addition, we have also shown that a group of EVRs in the stony corals cluster within the environmental PLV clade (discussed in the phylogenetics section), members that are known to exist as free virus particles in aquatic ecosystems^48^. Taken together, our findings provide compelling evidence that polintovirus and PLV-like EVRs within stony corals produce free viruses. However, we acknowledge that direct, microscopic observation of free viruses originating from endogenous polintoviruses is yet to be made^43, 51^ and future targeted studies will be required to definitively address this question.

## Conclusion

The discovery of a large diversity of polintoviruses and PLV relatives in stony coral genomes and their viromes lead to a number of questions regarding their roles in the ecology, physiology, and genome evolution of this important group of metazoans. The complex evolutionary histories and possible multiple origins of these endogenous elements within the stony coral genomes suggest that they may have originated from previous infections and propagated through vertical transfers and within-genome paralogous expansions. However, future studies focusing on active infections and identification of free polintovirus particles specific to corals will be instrumental in clarifying their origins and integration mechanisms. As these elements seem to have gone through massive lineage-specific expansions in several coral species, our results pose questions regarding their physiological effect and potential for promoting genetic diversity in corals. Particularly relevant is whether the dynamics of these elements could shape the environmental adaptation and response to diverse stressors in stony corals in the global ocean.

Addressing the questions related to the ecological and evolutionary importance of polintoviruses and molecular aspects of their interactions with diverse organisms will require a range of experimental approaches. A key future direction that emerges from our observation is the need to decipher the replication strategy of polintoviruses. Multiple studies have suggested that polintoviruses likely exist as free virus particles^48, 51, 73^, although direct observation of the particles is yet to be made^50^. A carefully curated clonal replicate experiment at different life-stages of the organism hosting these elements, along with multi-omic analysis at different growth stages in response to diverse stressors might clarify the replication strategy of these elements. In addition, microscopic observation of free virus particles coupled with targeted gene-knockout approaches would substantiate these observations. Together, the outcome of such experiments could lead to a better understanding of the molecular events underpinning the ecological dynamics of polintoviruses.

Another possible direction is to perform similar genomic analysis of the members of the broader cnidaria phylum using better quality genome assemblies through long read technology. This could help us understand the evolutionary role of these highly prevalent endogenous viral elements within this important group of animals crucial for ocean ecosystem health. Resolving the genomic landscape of these elements through long read sequencing would help better quantify the level of expansion of these elements in certain lineages compared to others, along with resolving strain-level heterogeneity in expansion - which might be useful to understand the role of these elements in niche adaptation and species-specific variations in genome evolution.

Polintoviruses and related elements represent an understudied, yet pervasive driver of genome evolution in diverse eukaryotes ranging from protists to metazoans. Stony corals are a widely studied group of marine animals for which the knowledge base is still being actively developed; given their importance in the stability of ocean ecosystems and response to global change^74, 75^. Given the widespread prevalence of polintoviruses across the phylogeny of stony corals, they could also serve as important model systems for studying the ecological roles of these viruses and molecular mechanisms that govern their interaction with host organisms.

## Materials and Methods

### Creation of HMM profiles

All proteins identified as belonging to the *Adintoviridae* family of viruses were downloaded from NCBI^76^ (April 2022) and clustered to identify orthologous groups according to a previously reported approach^77^. Briefly, we created orthogroups of proteins using Proteinortho v.6.06^78^ with parameters: -e 1e-05 –identity 25 -p blastp+ –selfblast –cov 50 -sim 0.80. Proteins within each orthogroup were aligned using Clustal Omega^79^ and resulting alignments were trimmed using trimal^80^ (parameter -gt 0.1). The alignments were then converted to HMM profiles using the hmmsearch tool implemented in HMMER ^81^ (3.3.2; hmmer.org) with default parameters. Similar methods were used to construct HMM profiles from the protein clusters (PCs) generated from environmental polinton-like viruses (PLVs) reported by Bellas and Sommaruga (2021)^48^. In addition, we also used several additional HMM profiles specific to the MCPs of virophages and PLVs that were constructed and reported by Bellas and Sommaruga (2021) ^48^.

Polinton-like viruses are highly diverse and consequently their hallmark genes can show substantial sequence heterogeneity. Due to this, many of these genes cluster into distinct orthogroups^48^, however these orthogroups still all correspond to the hallmark genes. For our analyses related to identification of hallmark genes, we identified all the HMM profiles that correspond to the hallmark genes in the Bellas and Sommaruga (2021)^48^ data, based on the provided annotations along with the HMM profiles we generated from the hallmark genes of *Adintoviridae* members.

### Identification of Viral Regions

Utilizing the ViralRecall tool^82^ originally developed for the identification of endogenous giant virus signatures in eukaryotic genomes, we identified potential endogenous polintoviruses within 30 coral genomes retrieved from NCBI^76^ in April 2022 (representing the total number of genomes publicly available around that time). To this end, ViralRecall^82^ was tailored so it would identify candidate polintovirus, PLV and virophage-like regions, instead of giant viruses. This was facilitated by replacing the original GVOG HMM profile database that ViralRecall^82^ utilizes for the identification of giant virus regions with the HMM profiles generated using the protein clusters from the genomes reported in Bellas and Sommaruga (2021)^48^.

For annotation purposes, ViralRecall^82^ outputs the presence of virus-specific hallmark genes as part of the candidate genome identification. In our modified version of ViralRecall^82^, five hallmark genes present in polintoviruses, PLVs and virophages (MCP, FtsK ATPase, mCP, polB and integrase) are reported instead of the giant virus-specific hallmark genes.

This modified ViralRecall^82^ program was run on the contigs >10kb in length from each stony coral genomes with an e-value of <1e-05, a minimum viral region length of 10kb, and a minimum score of 1. A parameter was used (-fl) to include a 5kb overhang on each end of the identified viral candidates. All other parameters were set to default.

Resulting viral regions were filtered to exclude regions containing fewer than one MCP, one mCP, and one FtsK ATPase gene. (Cut-off established based on the genes that are commonly present in all known PLVs, polintoviruses and virophages^48, 51^). Length cut-off for the regions were then established at a minimum of 10kb and a maximum size of 50kb based on the average size of PLV genomes previously reported^39, 55^.

### Compilation of a MCP database for phylogenetic analysis

The ncldv_markersearch script^40^, originally developed to identify hallmark genes of giant viruses was modified to incorporate curated hallmark gene HMM profiles specific to PLVs, polintoviruses and virophages. After modification, ncldv_markersearch^40^ was run on all identified viral regions that met established cut offs. MCP hits obtained by this approach in each species then clustered at 90% sequence similarity cut-off utilizing CD-Hit^83^ (default parameters) to reduce complexity in the dataset while removing redundant sequences for downstream phylogenetic analysis.

Metavirome sequences were downloaded from data provided by Weynberg *et al*. (2017)^32^ and the fastq reads were quality-trimmed using BBduk^84^, and assembled using Megahit^85^ using default parameters. Proteins were predicted in the assembled contigs using prodigal with default parameters. The modified ncldv_markerserach program^40^ was then used to identify the MCPs in this dataset as described in the previous paragraph. MCP hits obtained in this manner were also clustered using a 90% similarity cut-off.

Bellas and Sommaruga (2021)^48^ compiled a dataset of genomes that contains PLV and virophages identified in their study, as well as PLV and virophage genomes from a number of previous studies and sources^53, 54, 61, 86, 87^. We used the data from this report as a comprehensive source of references for PLVs and virophages for our phylogenetic analysis. In addition to this, we also included MCPs that were previously reported for Adintovirus genomes^53, 87^, which we downloaded from NCBI. We created a final set of MCPs combining the MCP cluster representatives in our study (from stony coral EVRs and metavirome study), along with these aforementioned studies.

### Phylogenetic analysis of the MCPs

The compiled dataset of MCPs were aligned using MAFFT^91^, and the final alignment was quality trimmed using Trimal^80^ (parameter gt 0.1). A diagnostic phylogenetic tree using this alignment was then created using Fasttree^92^ which we investigated to manually remove any long branches. The remaining MCP candidates were re-aligned and a final maximum likelihood tree was constructed using IQ-Tree v.2^93^ (parameter -B 1000), with 1000 bootstrap iterations ^94^ (Fig. 2). The tree was annotated using iToL ^95^.

### Validation of taxonomic assignment of virophages

The taxonomy of the reference virophage genomes were further validated using a recently reported script that can discriminate between virophages and PLV genomes based on gene content similarity^61^.

### Gene Annotation

Functional annotation of the proteins from the candidate viral regions in stony coral genomes were performed using the VOG database^96^, the annotations provided by Bellas and Sommaruga (2021)^48^, and our own compiled *Adintoviridae* protein data. To this end, the HMM profiles from the VOG database^96^ as well as those we generated from the protein clusters provided by Bellas and Sommaruga (2021)^48^ and our own NCBI curation, were searched against the proteins from the candidate viral regions in stony coral genomes with an e-value cut-off of 1e-05.

### Identification of Terminal Inverted Repeats

Terminal Inverted Repeats (TIR) flanking the candidate viral regions were located using the IUPACpal program^97^. The TIR length of polintoviruses is not defined, and likely varies widely. For our analysis, minimum and maximum TIR sizes were set at 100 base pairs (bp) and 2500 bp, respectively, based on the TIR analysis results from Hackl *et al* (2021) ^25^ as a guide. Maximum allowable mismatches between the TIR regions were set to 2%, and the minimum size allowed for the loop was set at 15kbp (based upon the average size of a standard polintovirus). For IUPACpal^97^, the following parameters were used: -M 2500 -m 100 -g 9000.

### Statistics

All statistical comparisons were run on R version 4.2.2^98^ using the ‘stats’ ^98^ and ‘EnvStats’^99^ packages.

### Gene expression profiling of *Stylophora pistillata* polintoviruses

To determine the expression profile of genes encoded by polintoviruses in *S. pistillata*, we downloaded the raw read libraries representing the transcriptome of *S. pistillata* colonies from Rädecker N *et al*. (2021)^71^. This data represents five colonies of *S. pistillata* both before and after undergoing heat stress treatment. Raw reads were quality trimmed using trim_galore (Krueger, 2017) with default parameters and were mapped to the five core genes (PolB, mCP, MCP, Integrase, FtsK ATPase) of the 74 *S. pistillata* polintoviruses using CoverM ^100^ with a stringent (99% identity) similarity cut-off. RPKM (reads per kilobase pairs per million mapped reads) values were calculated for each of the genes across all ten samples (5 control and 5 heat-treated).

## Data availability statement

The stony coral genomes analyzed in this study are publicly available at NCBI^76^ repository (See Supplementary Table 1 for accession numbers). All the identified viral regions, proteins, functional annotations and raw phylogenetic tree of MCPs are provided in an open access Figshare repository (https://figshare.com/articles/dataset/Data_Repository/23620479). The same repository also contains a version of the ViralRecall^82^ program that is modified for identification of polintoviruses, PLVs and Virophages from genome/metagenome sequence data.

## Author contributions

DS and MM co-conceptualized the study. DS performed the majority of the data analysis, synthesis, and writing. ZF performed part of the graphical data analysis. MM supervised the research and secured funding.

## Acknowledgements

MM and DS are funded by the Rosenstiel School of Marine, Atmospheric, and Earth Science, University of Miami.

