## Supplementary Materials for "Widespread Occurrence and Diverse Origins of Polintoviruses Influence Lineage-specific Genome Dynamics in Stony Corals"

**for**

This file contains supplementary figures 1-16 and supplementary table 1.

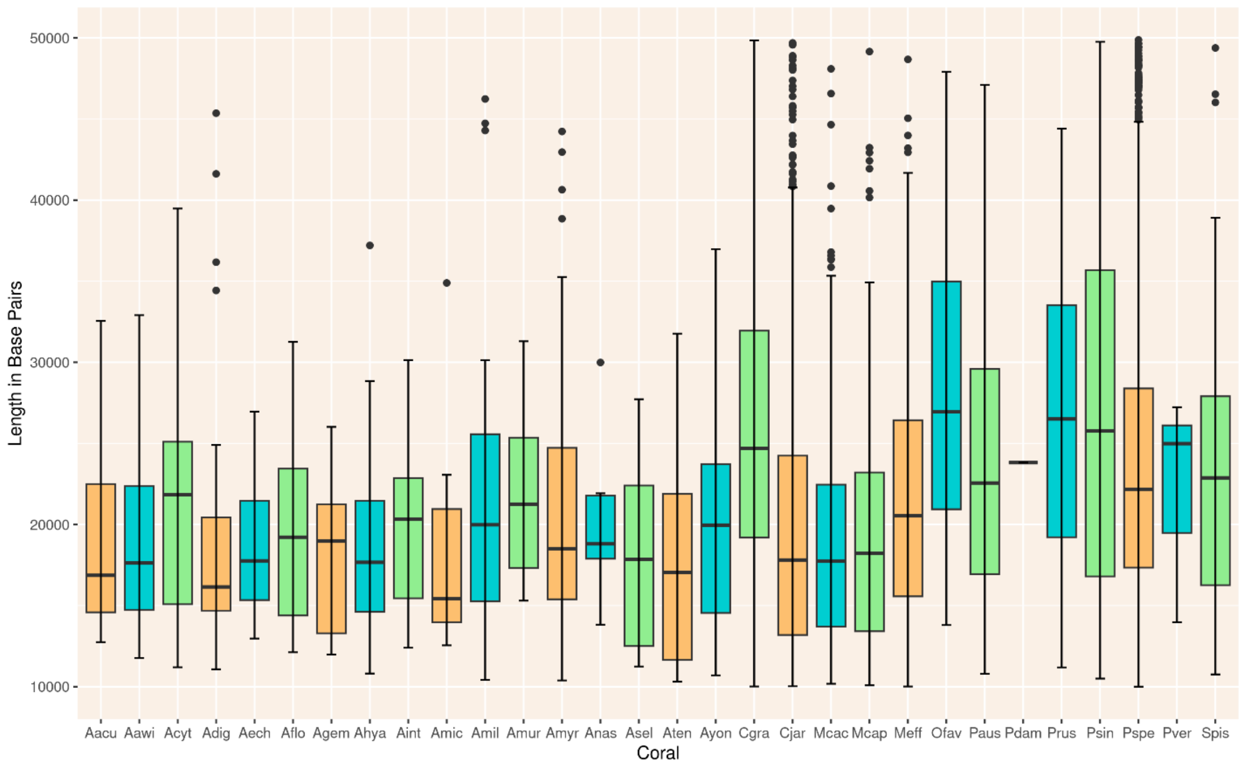


**Supplementary Fig.1:** Boxplot of viral region lengths by species. Figure generated using the ggplots package in R.

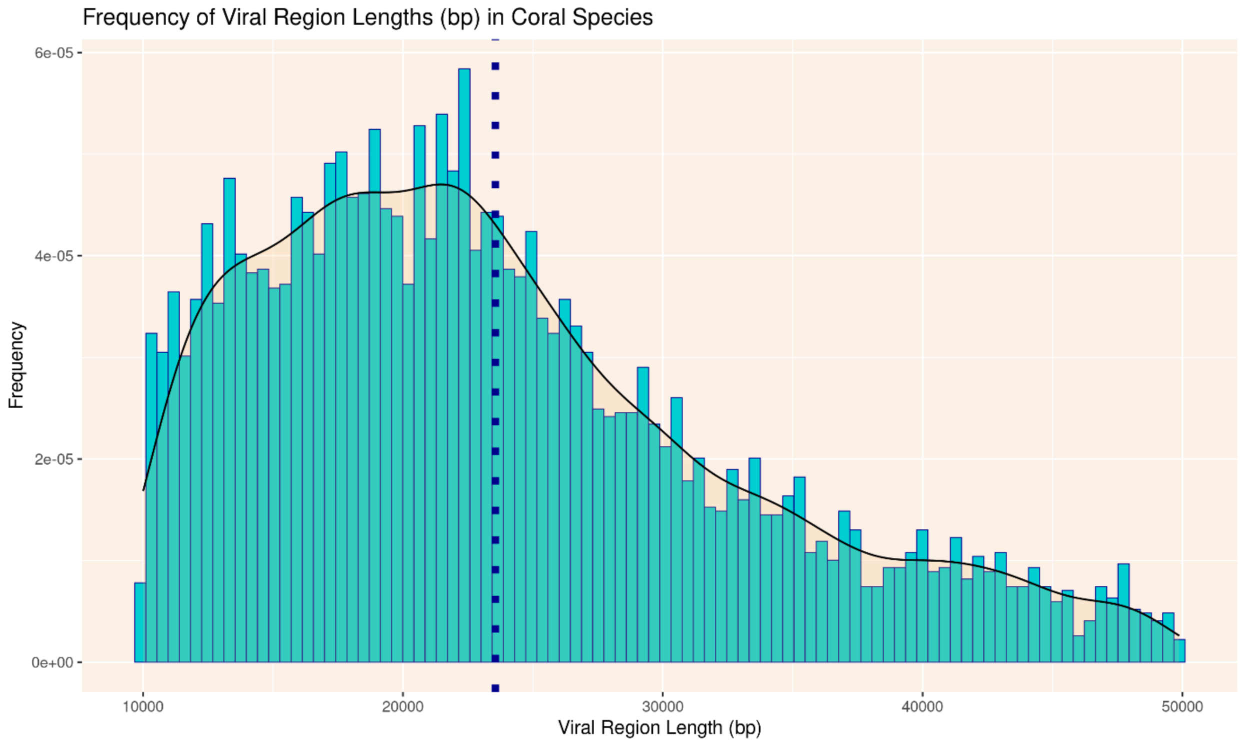


**Supplementary Fig.2:** Histogram of viral region length. Smoothing curve overlay. Dotted line shows median length. Figure generated using the ggplots package in R.

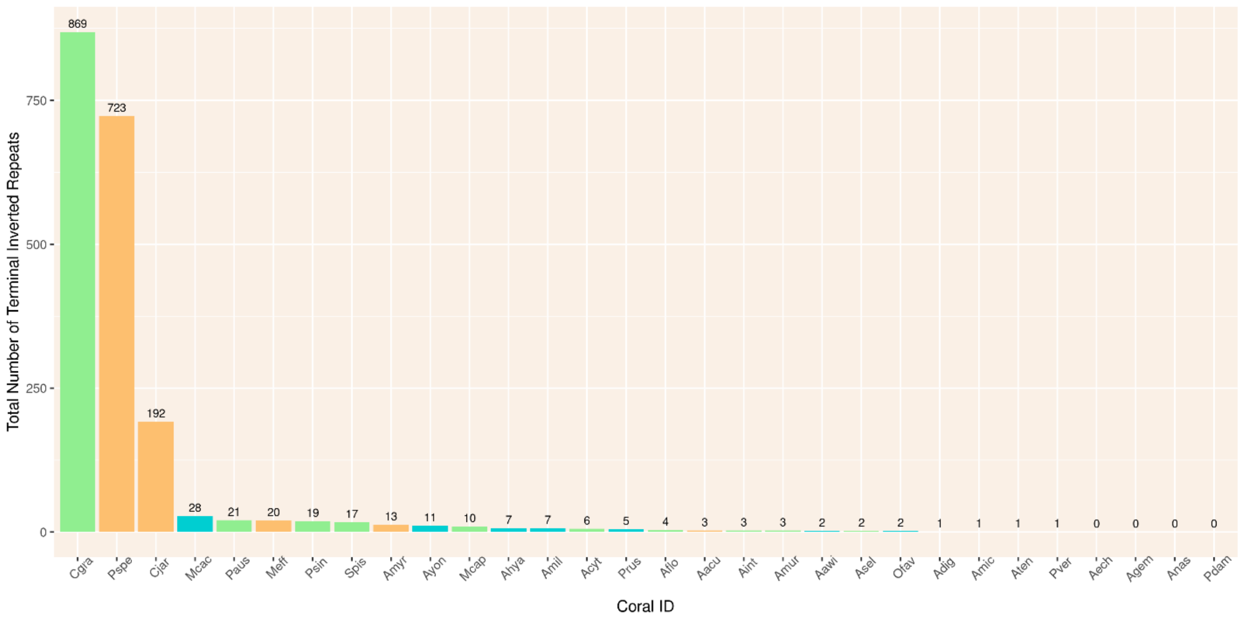


**Supplementary Fig.3:** Bar plot of total terminal inverted repeats by species. Figure generated using the ggplots package in R.

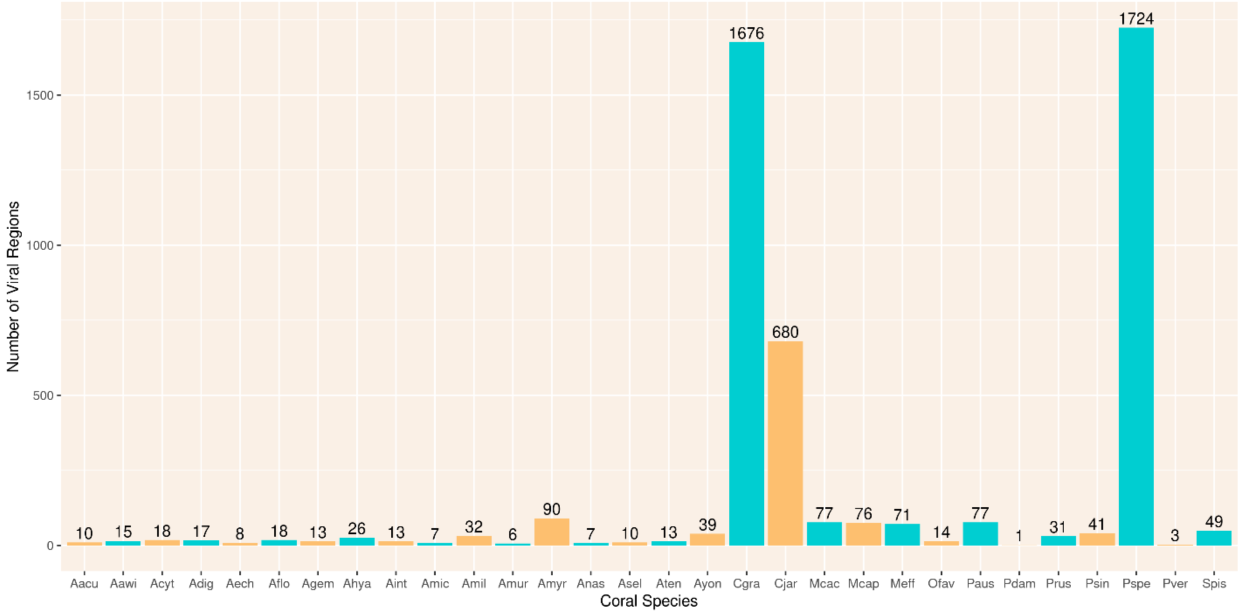


**Supplementary Fig.4:** Bar plot of total number of endogenized viral regions (EVRs) by species. Figure generated using the ggplots package in R.

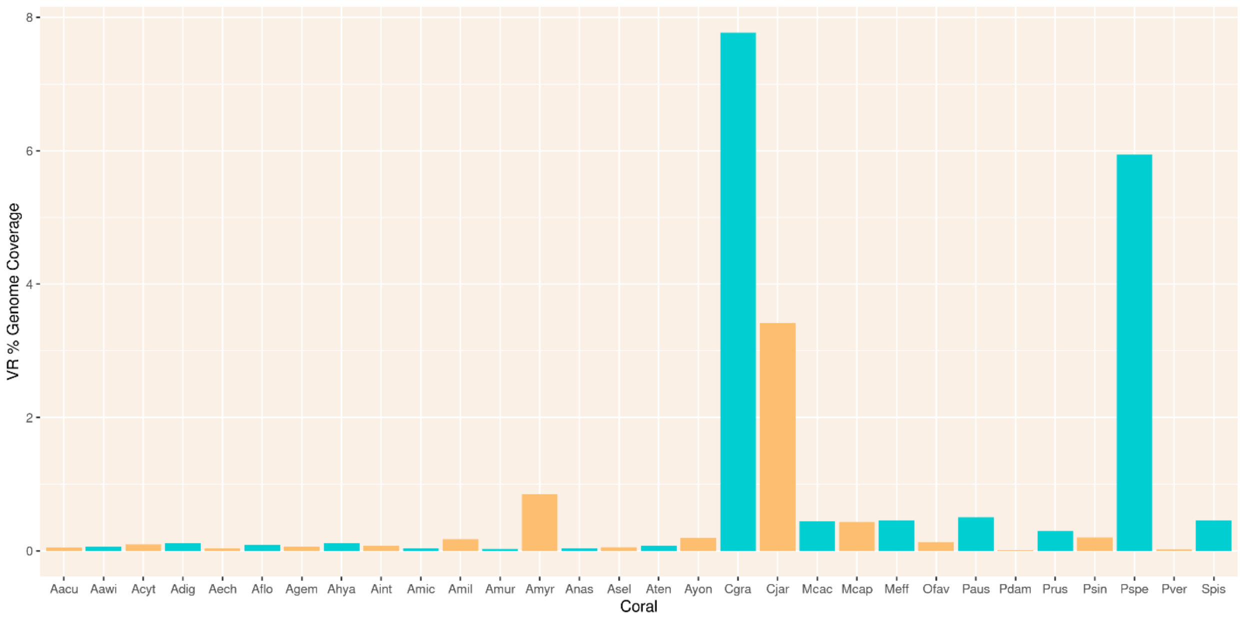


**Supplementary Fig.5:** Bar plot of percentage of coral genome length composed of endogenized polintovirus regions. Figure generated using the ggplots package in R.

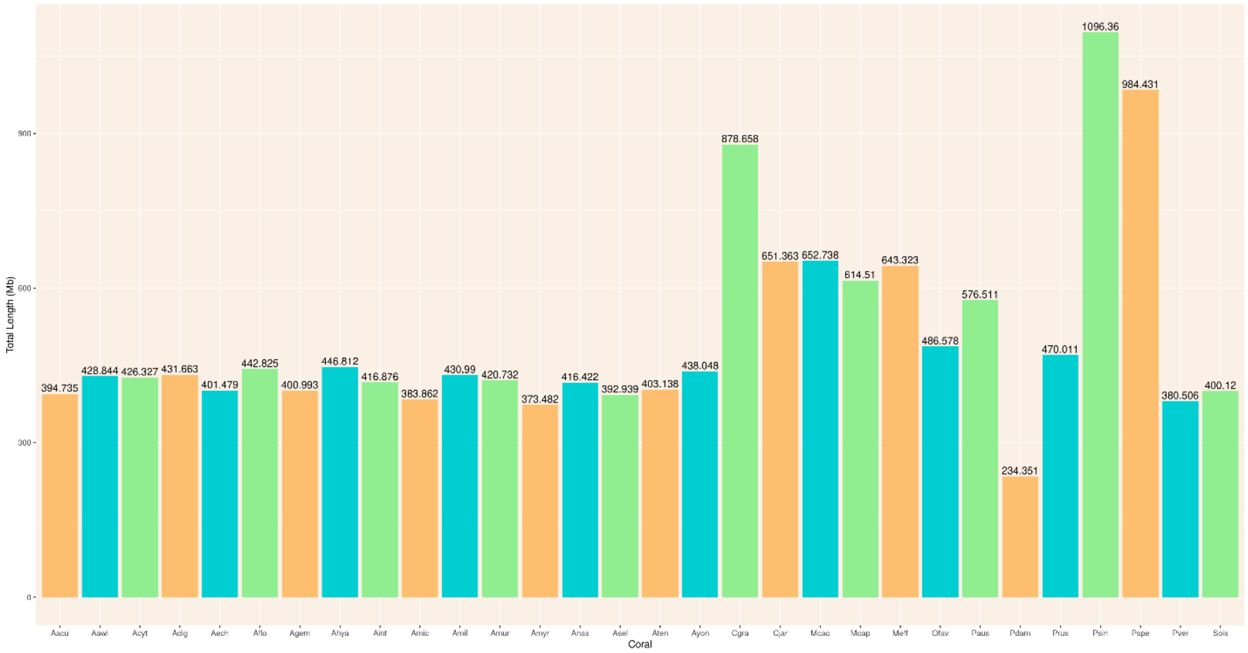


**Supplementary Fig.6:** Bar plot of coral genome length in mega-base pairs. Figure generated using the ggplots package in R.

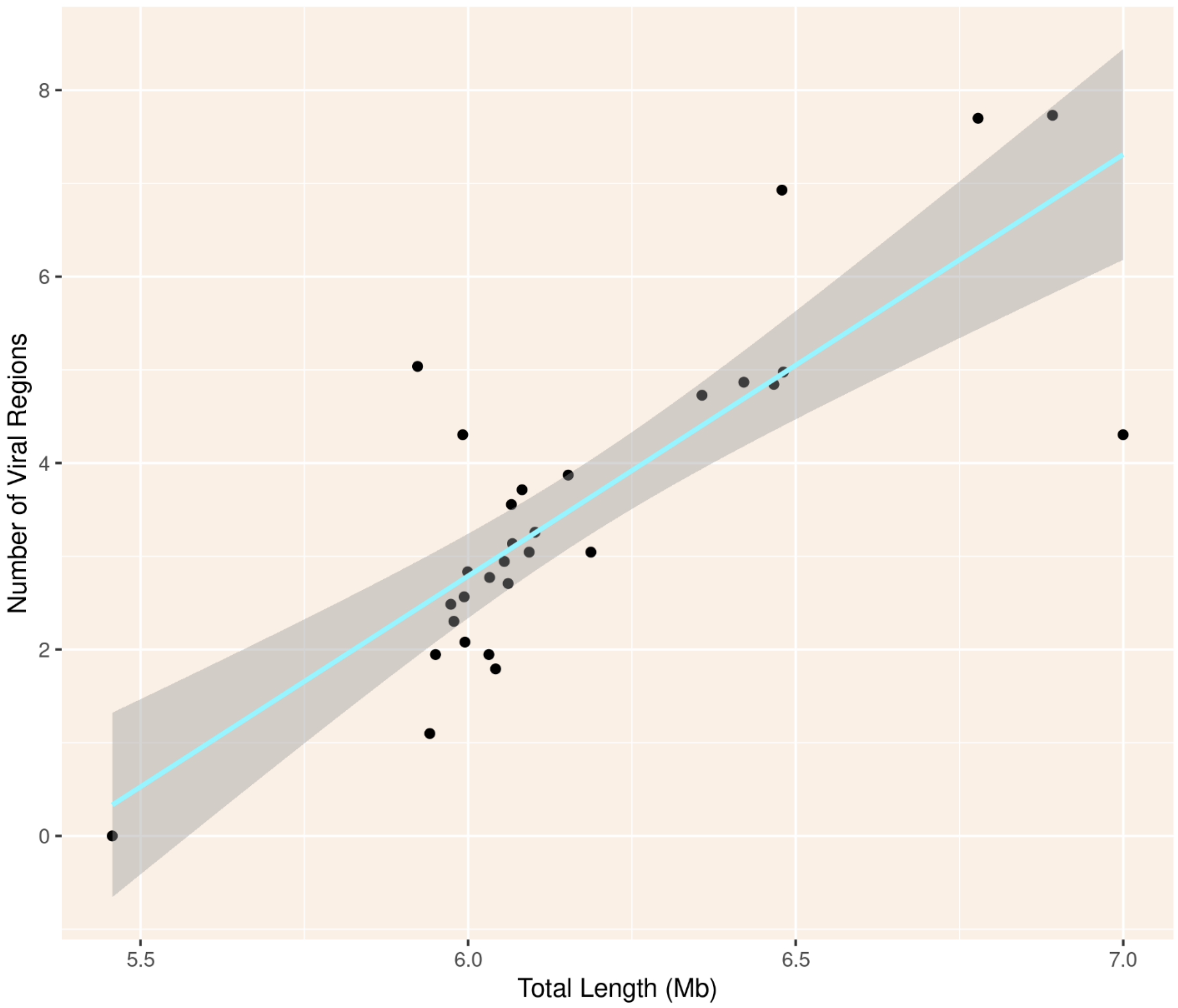


**Supplementary Fig.7:** Scatterplot of total number of endogenized viral regions in genome versus total length of genome. Blue line represents linear model overlay, and grey regions represent 95% confidence intervals. Figure generated using the ggplots package in R.

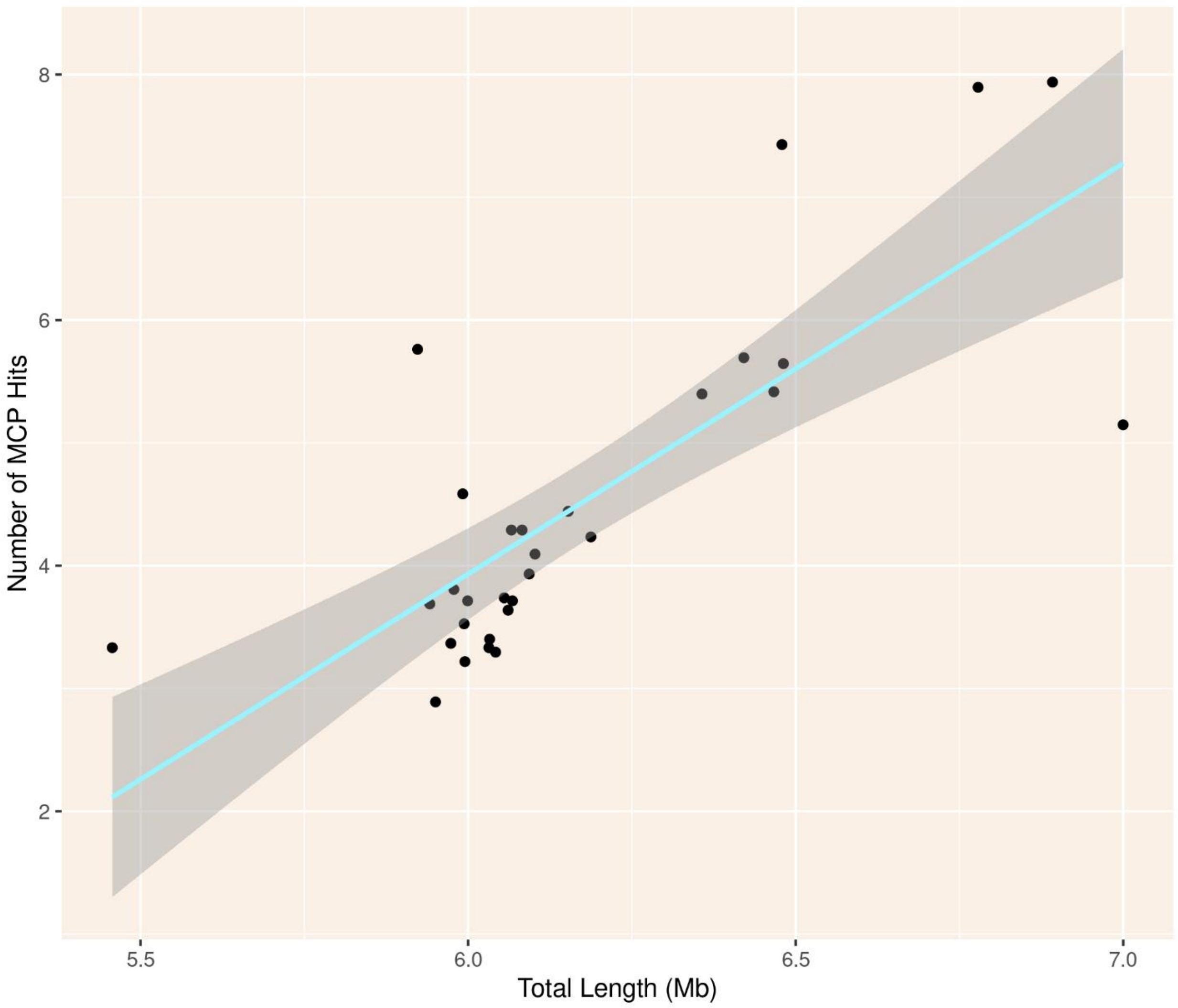


**Supplementary Fig.8:** Scatterplot of total number of MCP hits in genome versus total length of genome. Blue line represents linear model overlay, grey regions represent 95% confidence intervals. Figure generated using the ggplots package in R.

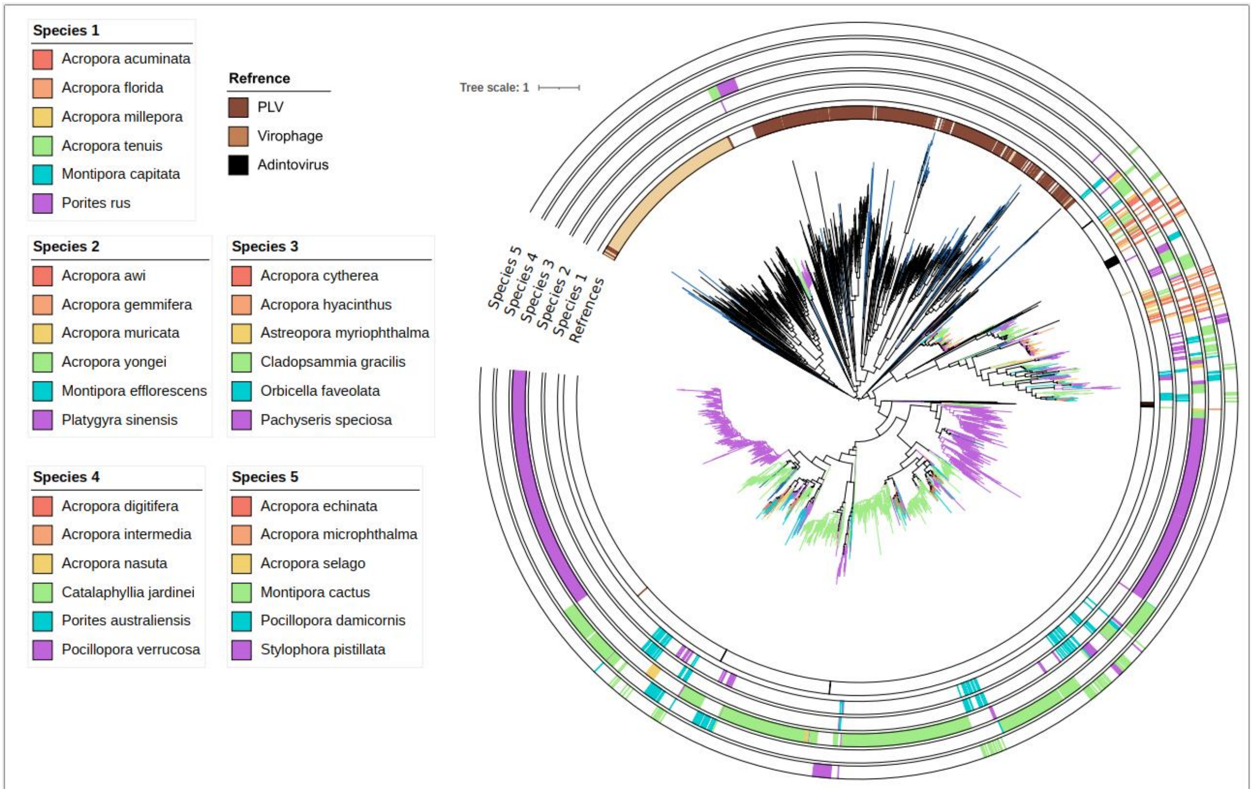


**Supplementary Fig.9:** Maximum likelihood phylogeny of polintoviruses in stony corals with each species separately annotated. As there are 30 distinct species, multiple annotation tracks were used for visual clarity, with each track showing data from six species. Tree generated utilizing IQ-Tree v.2.

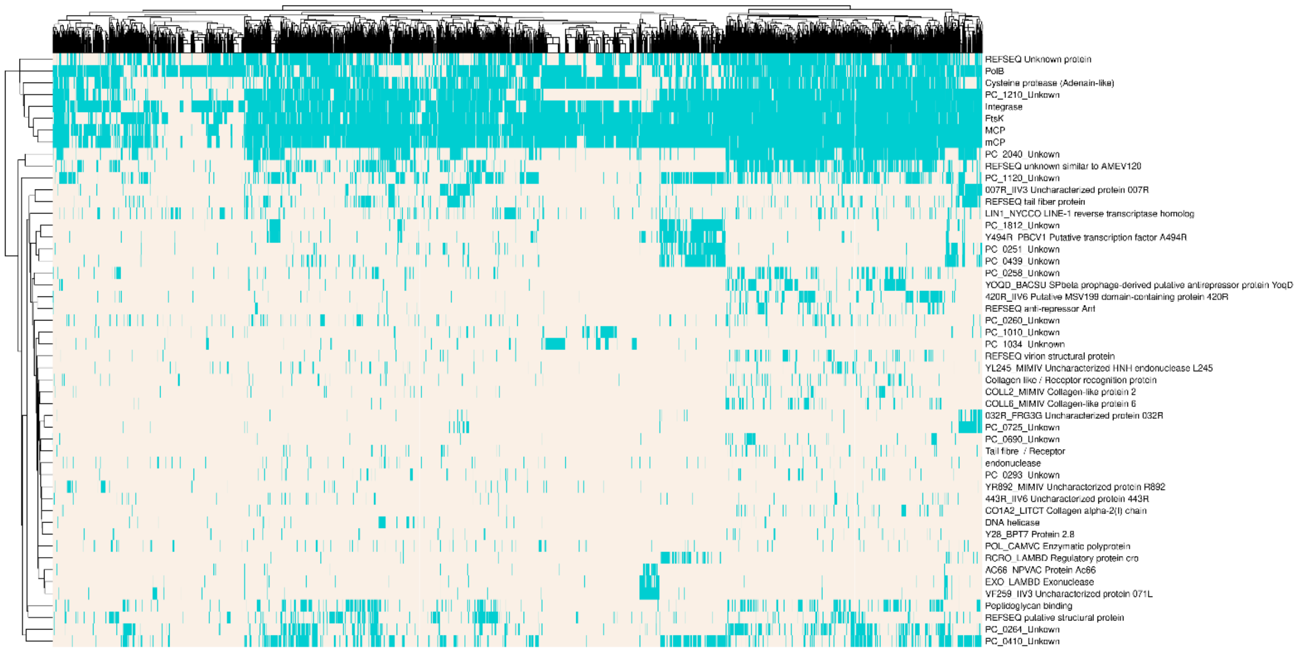


**Supplementary Fig.10:** Presence/absence heatmap of annotated polintovirus genes in each viral region. Annotated proteins were compiled with those occurring in at least 2% of total hits then run through the R package ‘pheatmap’.

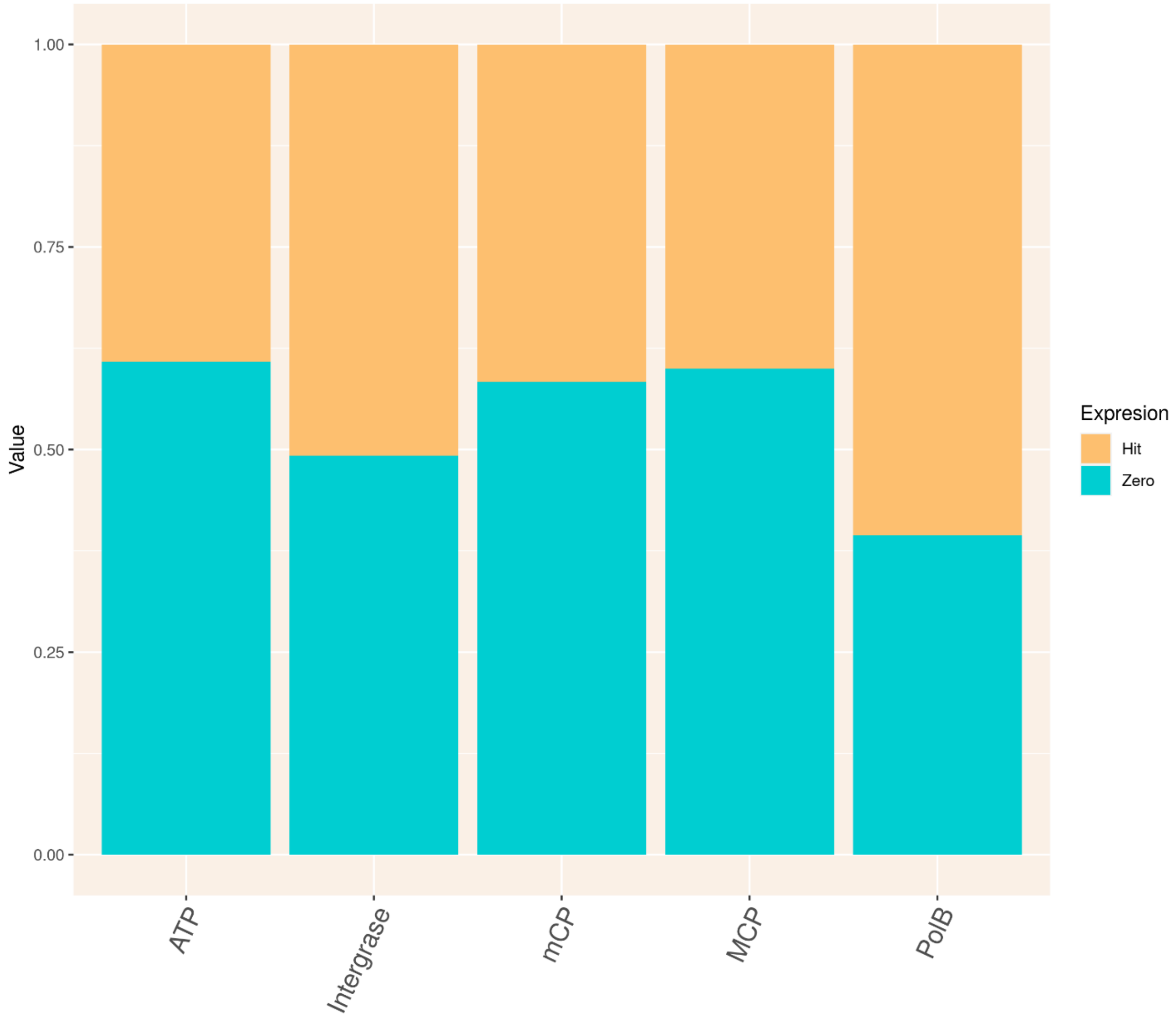


**Supplementary Fig.11:** Stacked bar plot of percentage of expressed core polintovirus genes from *S.pistillata* EVRs. Yellow represents % of EVRs where a particular gene is expressed, while blue represents % of EVRs without detectable expression of these genes. Figure generated using the ggplots package in R.

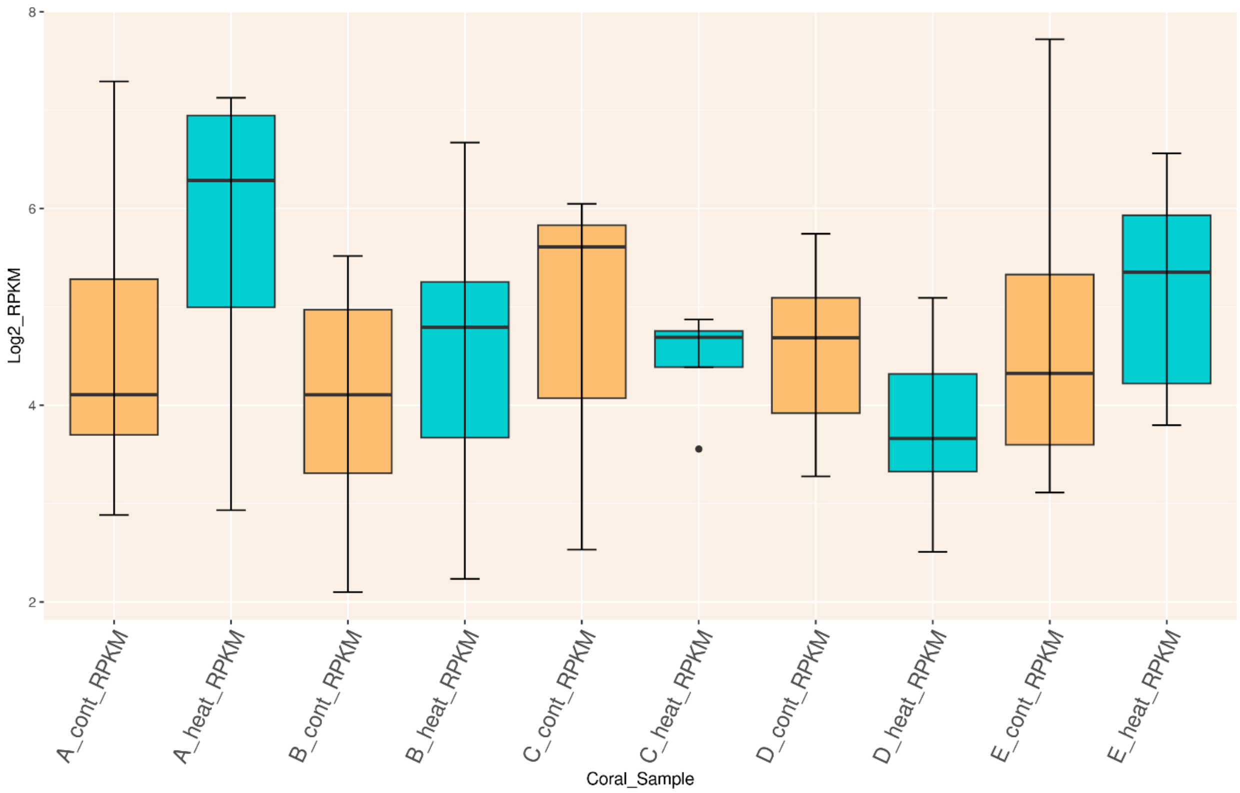


**Supplementary Fig.12:** Boxplot of distribution of major capsid protein (MCP) expression (RPKM) in 49 *S.pistillata* EVRs across different treatments (control and heat treated). Figure generated using the ggplots package in R.

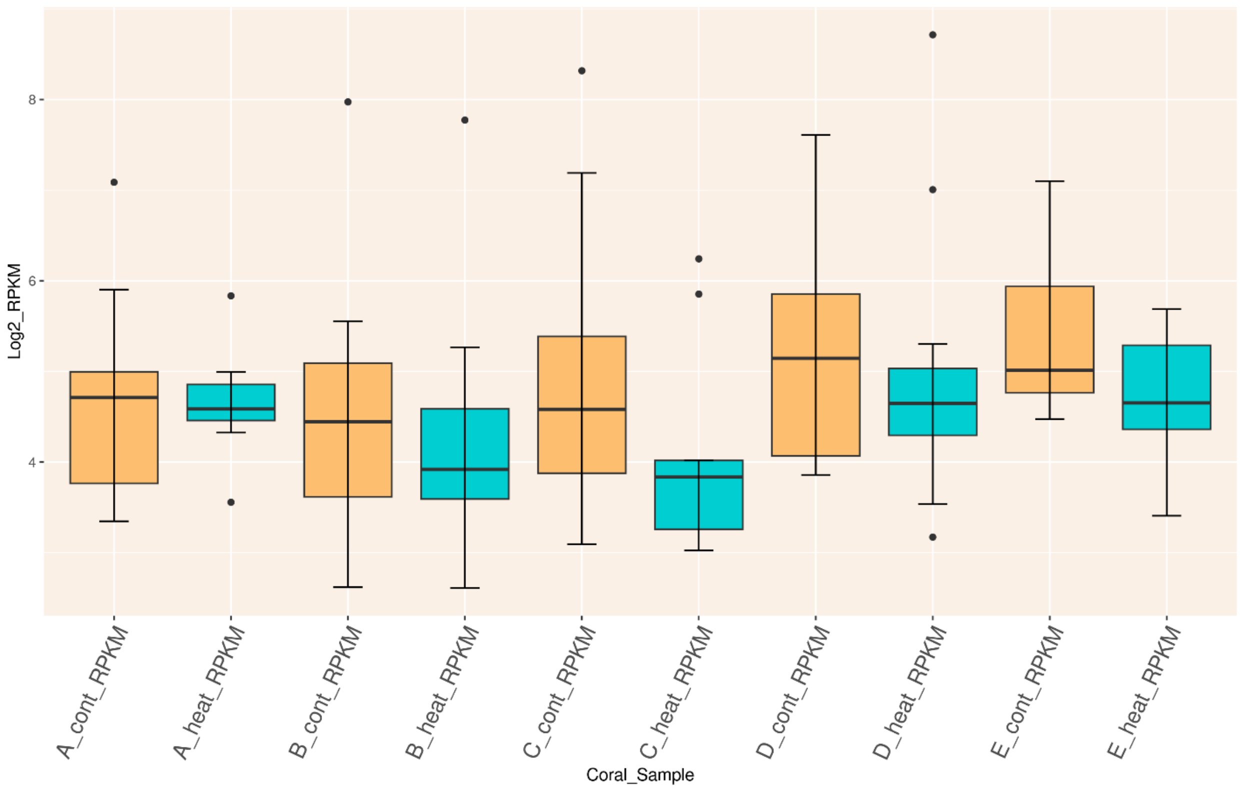


**Supplementary Fig.13:** Boxplot of distribution of minor capsid protein (mCP) expression (RPKM) in 49 *S.pistillata* EVRs across different treatments (control and heat treated). Figure generated using the ggplots package in R.

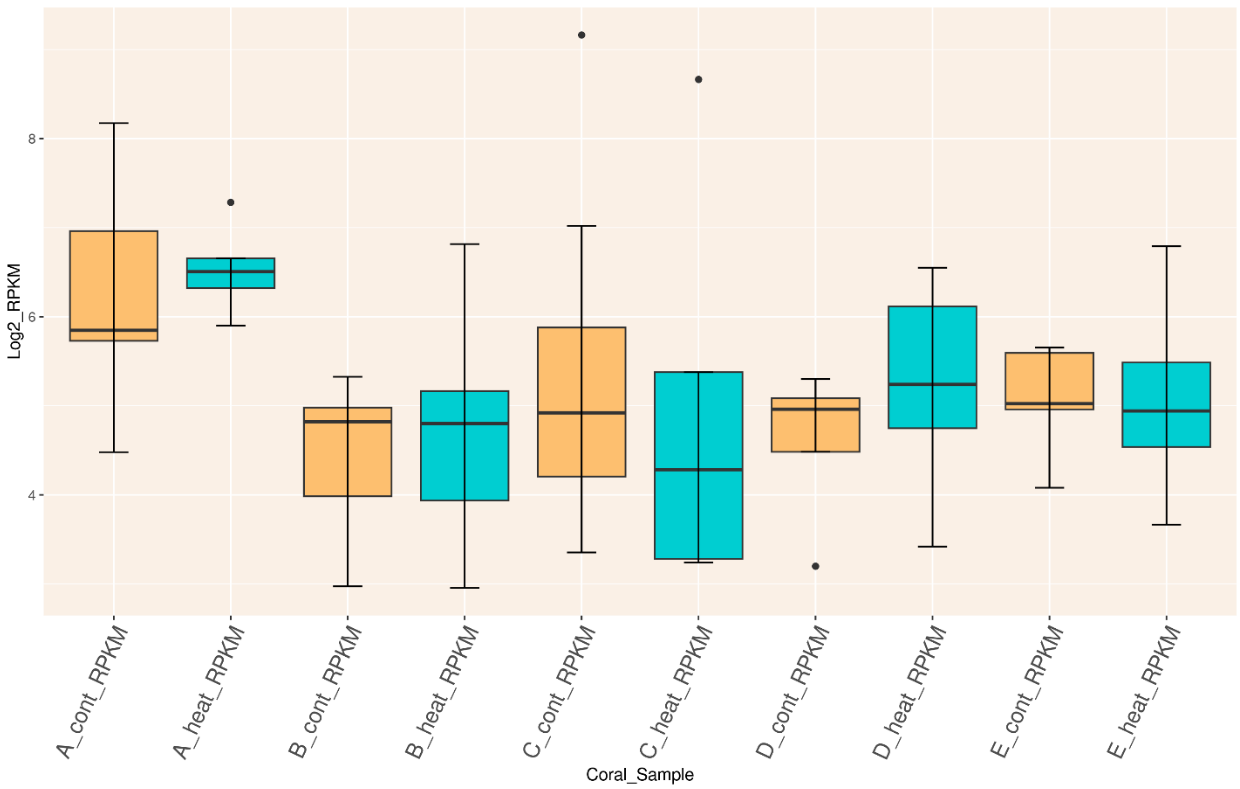


**Supplementary Fig.14:** Boxplot of distribution of ATPase protein expression (RPKM) in 49 *S.pistillata* EVRs across different treatments (control and heat treated). Figure generated using the ggplots package in R.

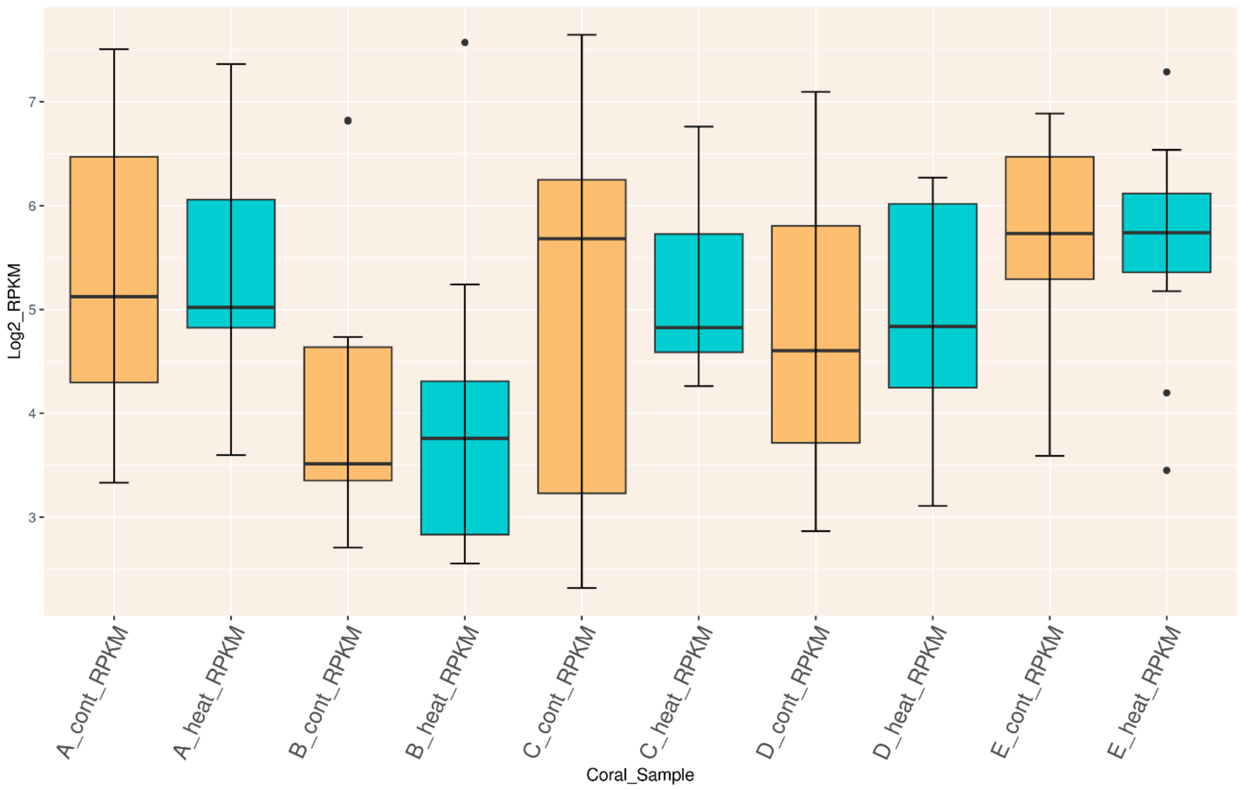


**Supplementary Fig.15:** Boxplot of distribution of Integrase protein expression (RPKM) in 49 *S.pistillata* EVRs across different treatments (control and heat treated). Figure generated using the ggplots package in R.

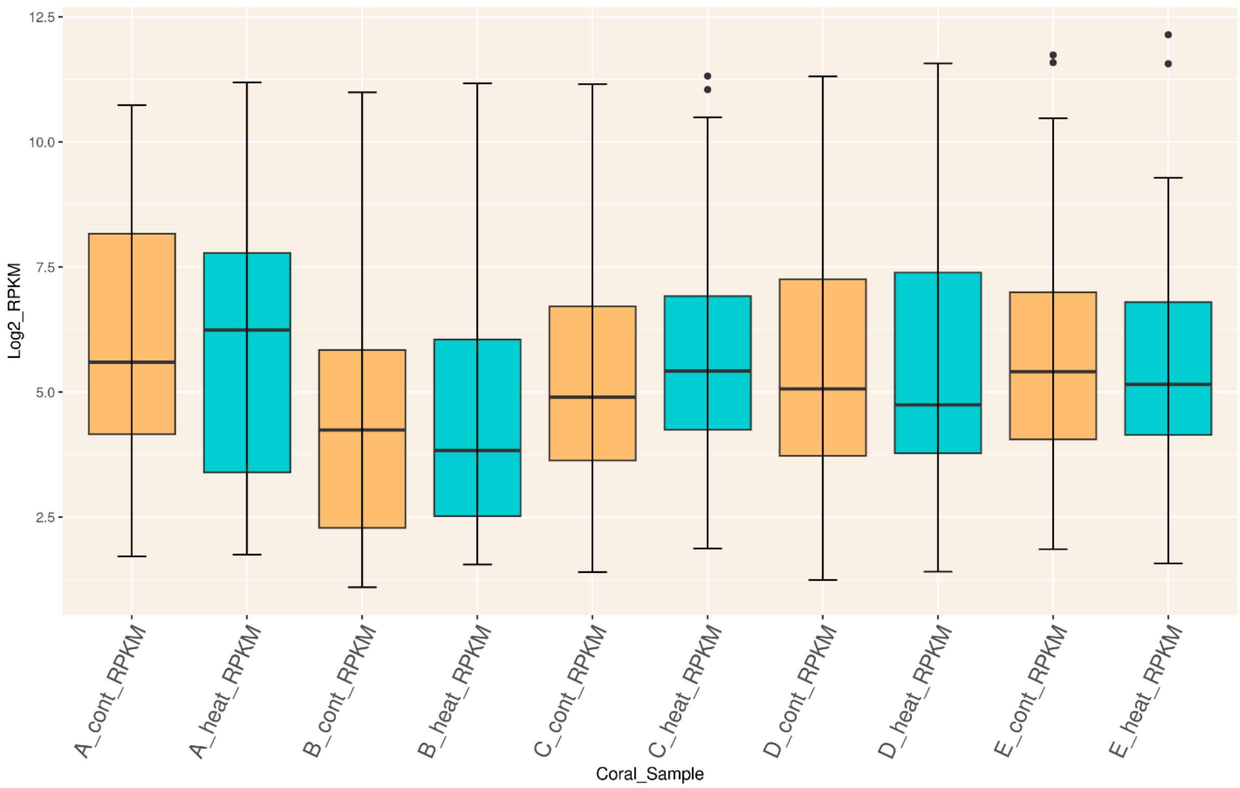


**Supplementary Fig.16:** Boxplot of distribution of DNA polymerase (polB) protein expression (RPKM) in 49 *S.pistillata* EVRs across different treatments (control and heat treated). Figure generated using the ggplots package in R.

**Supplementary Table.1:** NCBI accession numbers of the stony coral genomes use in this study.

| | Coral Spieces | Acseion Number |
|---|---|---|
| 1 | Catalaphyllia jardinei | JAEMPF000000000.2 |
| 2 | Cladopsammia gracilis | JAJGOT000000000.1 |
| 3 | Porites australiensis | BOPM00000000.1 |
| 4 | Platygyra sinensis | JAHPZR000000000.1 |
| 5 | Pachyseris speciosa | JAEMWC000000000.1 |
| 6 | Acropora intermedia | BLFH00000000.1 |
| 7 | Acropora selago | BLFM00000000.1 |
| 8 | Acropora microphthalma | BLFI00000000.1 |
| 9 | Acropora gemmifera | BLFF00000000.1 |
| 10 | Acropora echinata | BLFD00000000.1 |
| 11 | Acropora cytherea | BLFB00000000.1 |
| 12 | Acropora awi | BLFA00000000.1 |
| 13 | Acropora acuminata | BLEZ00000000.1 |
| 14 | Pocillopora verrucosa | AAVTL000000000.1 |
| 15 | Montipora capitata | RDEB00000000.1 |
| 16 | Montipora efflorescens | BLFP00000000.1 |
| 17 | Porites rus | OKRP00000000.1 |
| 18 | Astreopora myriophthalma | BLFK00000000.1 |
| 19 | Acropora muricata | BLFJ00000000.1 |
| 20 | Acropora yongei | BLFN00000000.1 |
| 21 | Acropora nasuta | BLFL00000000.1 |
| 22 | Acropora florida | BLFE00000000.1 |
| 23 | Pocillopora damicornis | RCHS00000000.1 |
| 24 | Acropora hyacinthus | GCA_020536085.1 |
| 25 | Orbicella faveolate | GCF_002042975.1 |
| 26 | Stylophora pistillata | LSMT00000000.1 |
| 27 | Acropora digitifera | GCF_000222465.1 |
| 28 | Montipora cactus | BLFO00000000.1 |
| 29 | Acropora tenuis | BLAZ00000000.1 |
| 30 | Acropora millepora | GCF_013753865.1 |
